# Allele-specific correction of dominant Best vitelliform macular dystrophy in patient-derived retinal pigment epithelium

**DOI:** 10.64898/2026.07.09.737567

**Authors:** Sushma Kalmodia, Jennifer G. Aparicio, Kayla Stepanian, Assaf Beck, Andrew Salas, Narine Harutyunyan, Patricia Galvan, Jinlun Bai, Maya Hayun, Meng Li, G. Esteban Fernandez, Mark W. Reid, Ryan J Schmidt, Catherine Argyriou, David Cobrinik, Aaron Nagiel

## Abstract

Autosomal dominant Best vitelliform macular dystrophy (BVMD) caused by variants in the *BEST1* gene is characterized by dysfunction of the macular retinal pigment epithelium (RPE) and secondary degeneration of the photoreceptors. There are currently no approved treatments for BVMD, and owing to its dominant nature, there remains uncertainty regarding the utility of traditional gene augmentation. Here we evaluated whether a dominant pathogenic *BEST1* allele can be corrected by base editing in differentiated RPE cells. We identified a patient with a likely pathogenic *BEST1* c.851A>G (p. Tyr284Cys) variant that was amenable to cytidine base editing. After establishing patient-derived induced pluripotent stem cells (iPSCs), we corrected the pathogenic variant in the iPSCs to obtain corrected iPSCs with the same genetic background. Corrected iPSC-derived RPE exhibited normalized monolayer appearance, improved barrier integrity, reduced cell death, and restored RPE-specific transcriptome. We then used a dual adeno-associated virus (AAV) split-intein system to deliver a CRISPR-associated protein 9 cytidine base editor (SpCas9-CBE) to *BEST1* c.851A>G mutant RPE monolayers and achieved editing of the pathogenic allele with a maximum efficiency of 13.42 ± 3.64% (mean ± SD). Together, these results demonstrate progress towards allele-specific base editing in a dominantly inherited retinal disorder.

## INTRODUCTION

Bestrophinopathies comprise a group of inherited retinal dystrophies (IRDs) characterized by retinal pigment epithelium (RPE)-photoreceptor complex dysfunction and degeneration. These are caused by pathogenic variants in the *BEST1* gene,^1–3^ which encodes the Bestrophin-1 (BEST1) protein expressed by RPE cells.^4–6^ This homopentameric calcium-activated chloride ion channel regulates intracellular calcium signaling, ion transport, and maintenance of the homeostatic milieu in the subretinal space.^7–9^ Pathogenic *BEST1* variants are associated with diverse molecular mechanisms, including loss-of-function, gain-of-function, and dominant-negative effects.^10^ More than 250 distinct pathogenic variants in *BEST1* have been identified, with about 90% resulting from dominant missense variants.^9,11,12^ Dominant variants in the *BEST1* gene lead to functional defects in the RPE, which result in the accumulation of vitelliform material and secondary photoreceptor degeneration in the macula, ultimately leading to vision loss.^13^ Autosomal dominant Best vitelliform macular dystrophy (BVMD) is the most common bestrophinopathy and occurs in about 1 in 60,000 individuals. The genetic and phenotypic heterogeneity of BVMD presents challenges for gene therapy development.^4,14^

BVMD has previously been modeled using induced pluripotent stem cell-derived retinal pigment epithelium (iPSC-RPE) from affected individuals.^10,15^ *BEST1* patient-derived iPSC-RPE exhibited reduced phagocytosis, increased apoptosis, and protein mislocalization, as well as defects in anion conductance.^15–18^ Previous studies using patient iPSC-derived RPE and canine disease models have demonstrated the therapeutic potential of *BEST1* gene augmentation for recessive bestrophinopathies, restoring Ca²⁺ channel activity in patient-derived models and improving visual function in a canine model.^19,20^ However, therapeutic efficacy in BVMD models appears to be largely limited to BEST1 variants affecting ion-binding domains.^21,18^ Thus, while BEST1 gene augmentation may provide therapeutic benefit in selected cases, it may not be effective for some dominant BEST1 variants.

Another approach for treating dominant BVMD is to selectively correct or silence the pathogenic allele without disrupting the normal allele using base editing.^11,22^ In preclinical animal models, base editing has been shown to correct pathogenic variants and restore gene function, demonstrating potential for treating retinal genetic disorders.^23,24^ However, to our knowledge, base editing approaches targeting BVMD have not yet been reported. In this study, we utilized patient-derived iPSC-RPE cells to model BVMD and test a base editing strategy delivering a cytosine base editor (CBE) via a split-intein dual AAVs.^25^

## RESULTS

### Identification of a BEST1 variant amenable to base editing

We identified a patient carrying a likely pathogenic *BEST1* c.851 A>G (p.Y284C) by focused exome sequencing and computationally predicted that this variant may be amenable to base editing. At 9 years of age, fundus autofluorescence imaging revealed hyperautofluorescent vitelliform lesions in the macula of both eyes (Figure 1A). Visual acuities were 20/40 in the right eye and 20/50 in the left eye. The *BEST1* variant was confirmed to be heterozygous by Sanger sequencing (Figure 1B). To correct the pathogenic allele, we developed a base editing strategy using a single-guide RNA (sgRNA) targeting the noncoding strand (minus strand) of the *BEST1* c.851A>G variant (Figure 1C). The editing was designed to convert a C (the cognate base of the variant G at cDNA position 851) to a T, reverting the c.851A>G variant to the wildtype A.

**Figure 1:**
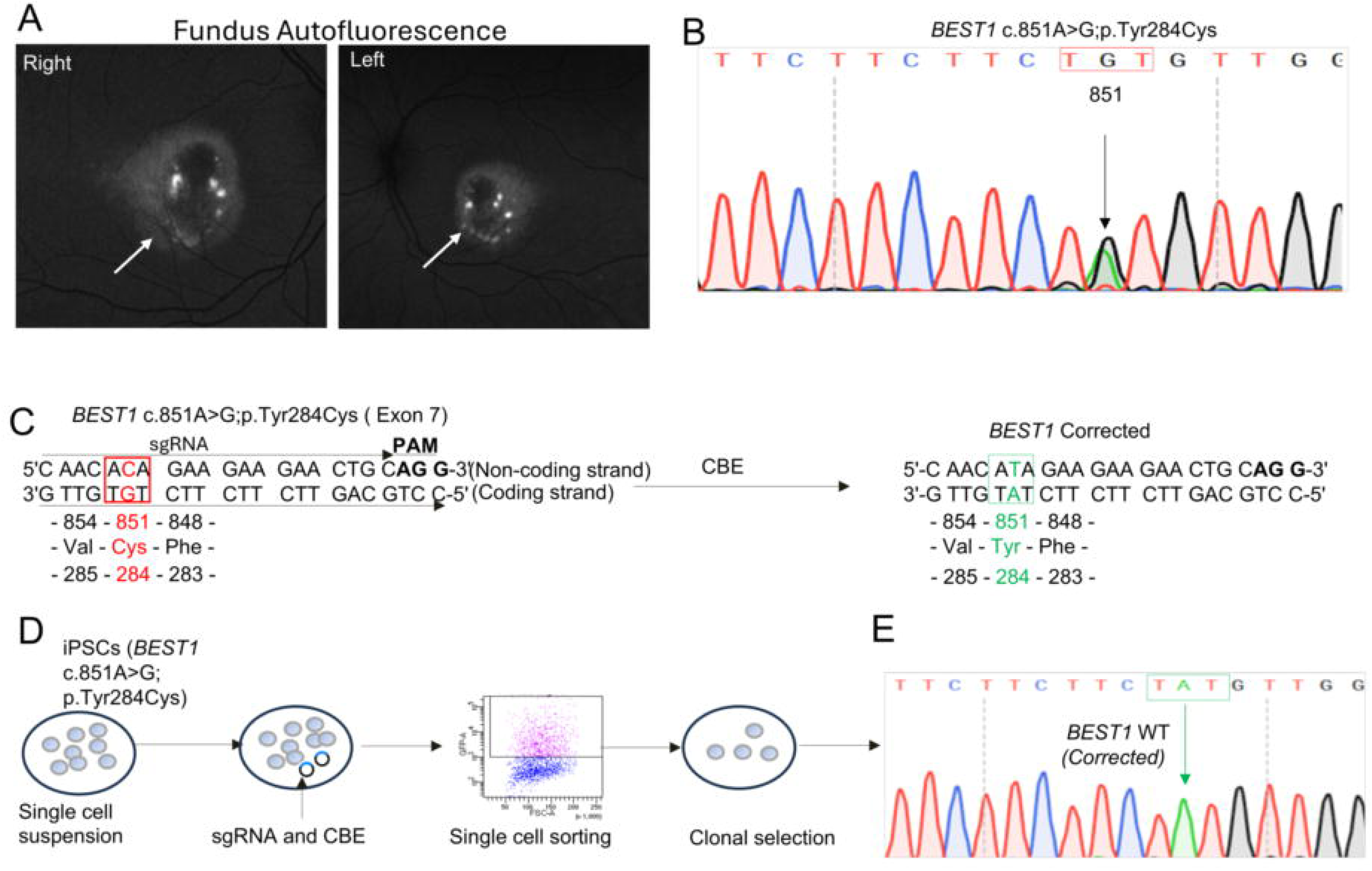
*BEST1* variant identification and gene editing strategy. (A) Fundus autofluorescence images of this Best disease subject harboring the *BEST1* c.851A>G (p.C284Y) variant. Hyperautofluorescent lesions are evident (white spots indicated by white arrow). (B) Genotyping (Sanger sequencing of PCR amplified genomic DNA) confirmed the *BEST1* variant in iPSC lines derived from the blood of *BEST1* subject. Heterozygosity at position 851 (black arrow) is evidenced by overlapping G (black) and A (green) peaks in the Sanger sequencing. Schematic diagram of *BEST1* cytidine base editing (CBE). CBE converts C (highlighted red) to T (highlighted green) in the target site (c.851) of the *BEST1 c.851A>G* variant, correcting amino acid cysteine (Cys) to tyrosine (Tyr). Protospacer sequence (sgRNA), and PAM (NGG; shown in **bold**) are labelled on the non-coding strand. (D) Workflow to create an isogenic control line. (E) Sanger sequencing confirming *BEST1* c.851 correction in the corrected iPSC lines.

### Generation of patient-derived iPSCs and isogenic control lines

To determine whether the *BEST1* c.851A>G variant causes RPE dysfunction, we established clonal iPSC lines from patient erythrocyte progenitor cells isolated from peripheral blood (Supplementary Figures S1A-S1F). We then created multiple isogenic control lines from two independent iPSC clones carrying the *BEST1* c.851A>G variant by co-transfecting with the *BEST1* targeted sgRNA and CBE4-SpG (P2A–EGFP) plasmid (Figure 1D and Supplementary Figures S2A-S2D). Cells transiently expressing EGFP were sorted, single-cell cloned and genotyped by amplifying the affected exon. Edited clones carrying the corrected C >T substitution and lacking off-target edits (Figure 1E and Supplementary Figures S3A - S3C) were selected for functional studies.

### Characterization of BEST1 variant-associated phenotypes

*BEST1* c.851 A> G mutant and *BEST1* corrected iPSC lines were differentiated into RPE cells using a previously established protocol with modifications (Supplementary Figure S4A).^26^ Hereafter, iPSC-derived RPE cells carrying the *BEST1* c.851A>G variant are referred to as ‘*BEST1* mutant RPE’, and RPE cells derived from *BEST1-*corrected iPSC lines are referred to as ‘corrected RPE’. Both *BEST1* mutant RPE and corrected RPE formed monolayers with cobblestone-like packing of pigmented epithelium, which is characteristic of well-differentiated RPE cultures (Figure 2A and 2B).^27^ However, *BEST1* mutant RPE exhibited focal areas of cell loss over time compared with corrected *RPE* (Figure 2A and Supplementary Figure S4B).

**Figure 2:**
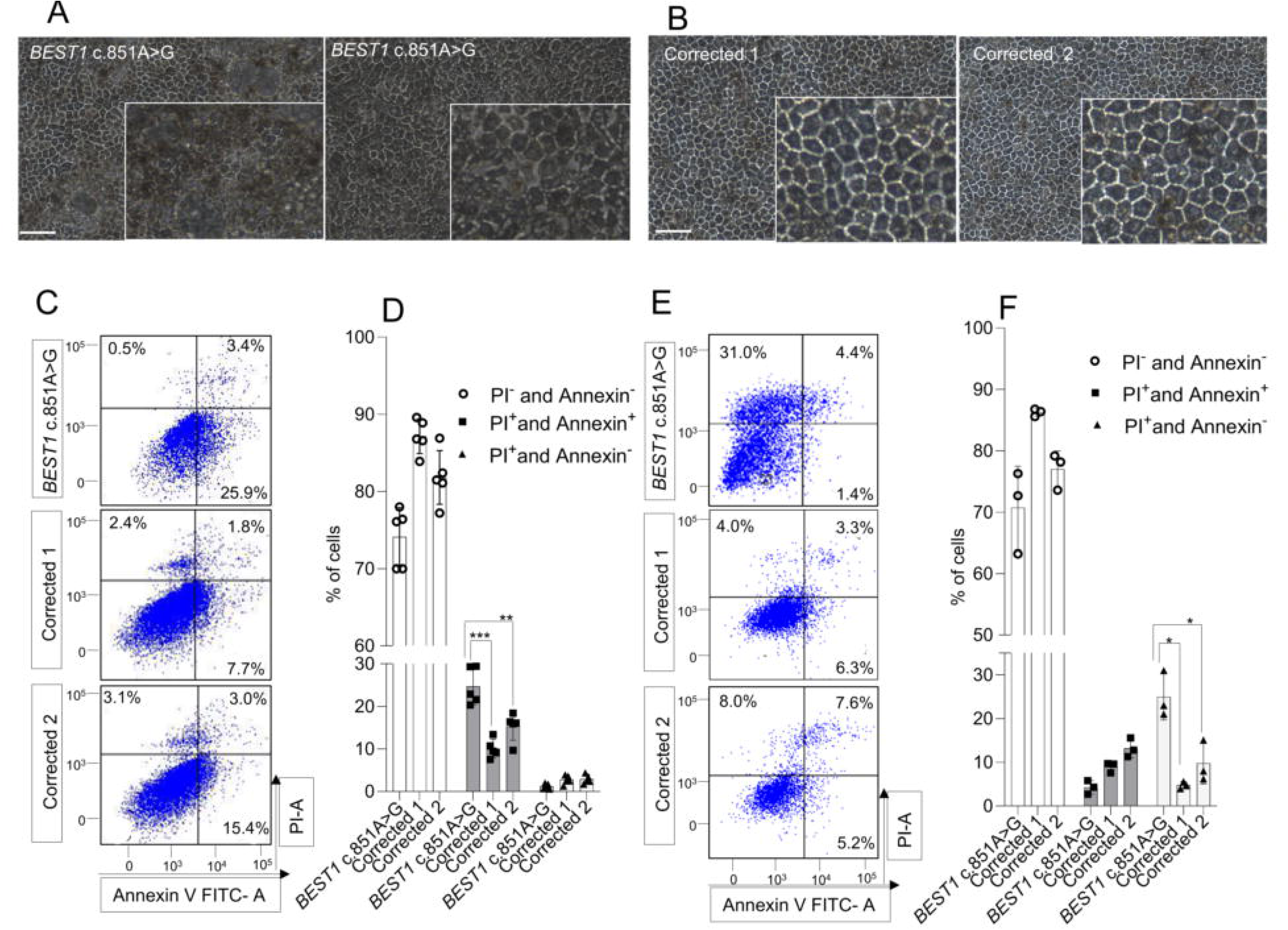
Morphological and cellular alterations in RPE derived from *BEST1 c.851A>G* and corrected clones. (A and B) Light microscope images of RPE cells at 10 weeks of differentiation. (A) *BEST1* c.851A>G RPE cells exhibit irregular morphology and regions of cell loss. (B) Corrected clones exhibit characteristic polygonal morphology and uniform monolayers. Inset represents a zoomed-in region highlighting the difference between mutant and corrected RPE. Images are representative of ≥4 independent RPE differentiation experiments. Scale bar: 20 μm. (C-F) Flow cytometry analysis of RPE cell population at 4 weeks (C-D) and 8 weeks (E-F) (post passage 1) using Annexin V-FITC and Propidium Iodide (PI) staining. This analysis distinguishes cell populations as follows: viable cells (Annexin V⁻/PI⁻), early apoptotic cells (Annexin V⁺/PI⁻), late apoptotic cells (Annexin V⁺/PI⁺), and necrotic cells (Annexin V⁻/PI⁺). Data are representative of 3–5 independent replicates per condition, from 3 independent experiments. Apoptotic cell population represents the combined early and late apoptotic cells. * Indicates a significant difference between *BEST1* c.851A>G mutant and corrected clones in apoptosis at 4 weeks and necrosis at 8 weeks (*p<0.05 and ** p<0.01,***p<0.0001)

To elucidate the potential cause of cell loss in *BEST1* mutant RPE, we evaluated cell death in RPE cultures using Annexin V–FITC and Propidium Iodide (PI) staining, which indicates the proportion of early apoptotic (Annexin-V^+^, PI^−^), late apoptotic (Annexin-V^+^, PI^+^), and necrotic (Annexin-V^−^, PI^+^) cells. At four weeks of culture, the total proportion of apoptotic cells was ∼ 25% in *BEST1* mutant lines but only ∼ 10-15% in corrected clones (p<0.005) (Figures 2C and 2D). In contrast, the proportion of exclusively PI positive cells, representing necrosis, was at most 2–3% of the total cell populations. At eight weeks, however, the proportion of necrotic cells rose to ∼ 25% in *BEST1* mutant RPE but was only 5-10% in the corrected clones (p=0.02) (Figures 2E and 2F). Collectively, these findings indicate that the *BEST1* mutation is associated with progressive RPE cell death, consistent with a previous study.^16^

To further investigate the molecular basis of BVMD associated with the *BEST*1 c.851A>G variant, we performed bulk RNA sequencing (RNA-seq) and differential gene expression (DGE) analysis on *BEST1* mutant and corrected RPE cells. We identified 1,975 upregulated and 1,292 downregulated differentially expressed (DE) genes in *BEST1* mutant RPE versus corrected RPE clone 1, and 1,978 upregulated and 1,404 downregulated DE genes in *BEST1* mutant RPE versus corrected RPE clone 2 (Figures 3A and 3B). Several canonical RPE genes were downregulated in the mutant RPE, including genes associated with extracellular structure organization (*CST3*), the visual cycle (*RGR, BEST1, RDH5, RPE65*), and RPE-enriched secreted proteins (*SERPINE3* and *SFRP5*) (Figure 3C). *SERPINE3* targets caspases involved in apoptosis, while *SFRP5* modulates Wnt signaling and is highly expressed in mature RPE. Among macula-specific DE genes (Figure 3D), *SULF1* and *WFDC1* were significantly downregulated in mutant RPE cells, consistent with previously reported roles *in BEST1* associated disease pathogenesis (Figure 3E).^28,29,30^ Our analysis also revealed changes in the solute carrier (SLC) and transmembrane (TMEM) family of genes, which are involved in metabolite and ion transport and fluid balance in the RPE. Notably, *TMEM97* (Sigma-2 receptor) and *TMEM163* are upregulated in *BEST1* mutant RPE. *TMEM97* is critically involved in regulating oxidative stress, apoptosis, cytotoxicity, and Ca2^+^ and K^+^ signaling, whereas *TMEM163* induces zinc efflux and is involved in zinc homeostasis, respectively (Figures 3E and 3F). *TMEM135*, which is downregulated in mutant RPE cells and has been implicated in macular degeneration, functions as a key regulator of lipid metabolism and mitochondrial dynamics among the SLC family, *SLC22A8* (a sodium-independent transporter) and *SLC6A13* (*GAT2*; a sodium- and chloride-dependent GABA transporter), were downregulated in *BEST1* mutant RPE cells. Thus, gene sets associated with phagocytosis, the visual cycle, pigmentation, macular RPE identity, and ion and fluid balance were differentially expressed due to the *BEST1* c.851 A>G variant (Supplementary Figures S5A-S5D).

**Figure 3:**
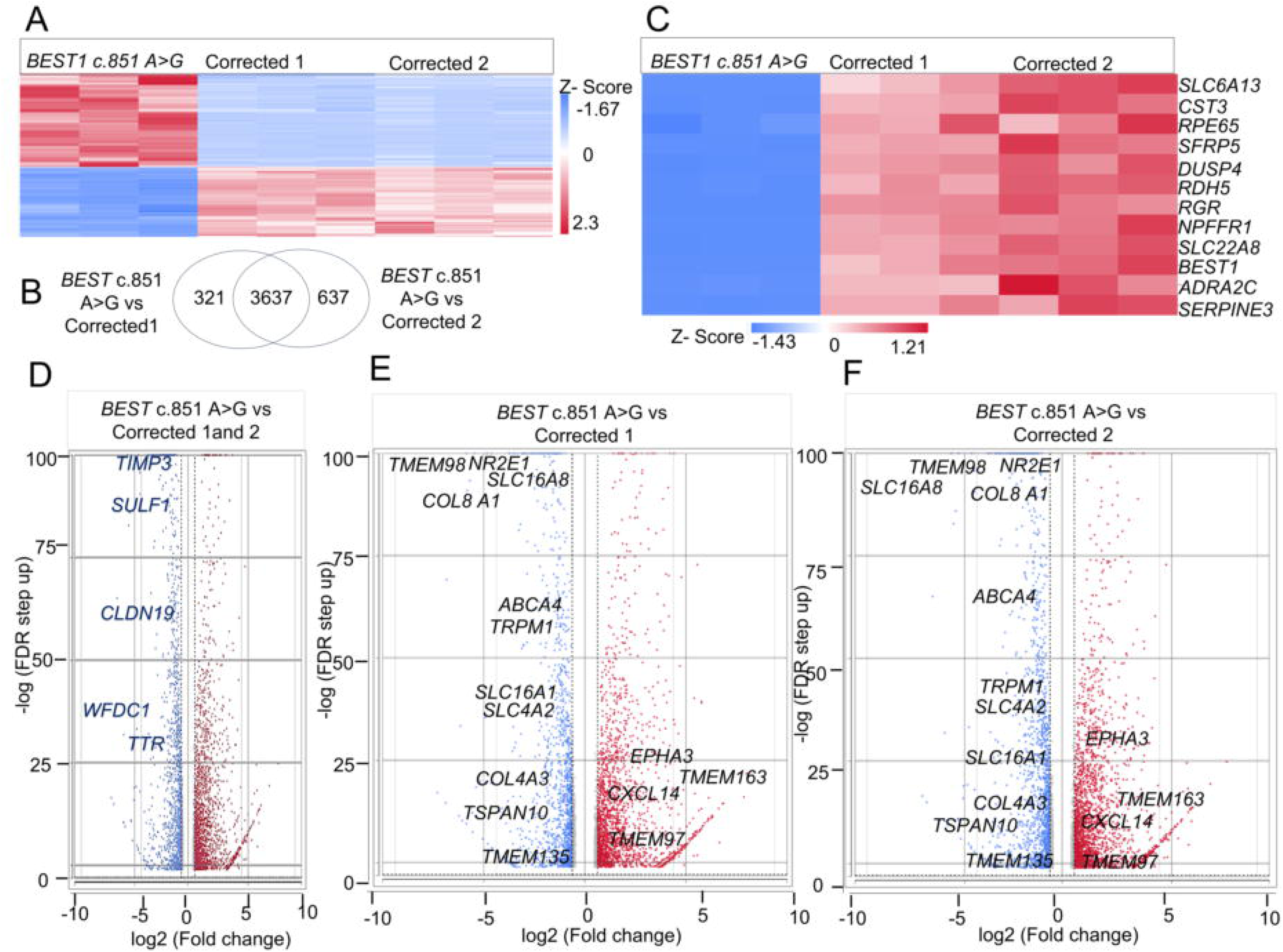
Transcriptome comparison of RPE cells of the *BEST1* c.851A>G variant and corrected clones. (A) Heat map of hierarchical cluster analysis (HCA) showing normalized gene expression counts across replicates for *BEST1* c.851 A>G, corrected clone 1 and corrected clone 2 iPSC-derived RPE cell lines (n=3 each). Corrected clones clustered together and showed expression patterns similar to each other and distinct from the *BEST1* c.851 A>G line. (B) Overlap of differentially expressed genes (DEGs) in *BEST1* mutant and corrected RPE clones. Venn diagram shows the total number of unique and overlapping significantly DEGs between *BEST1* c.851A>G and two corrected 1 and 2. DEGs were identified and filtered using the DESeq2 method (fold change ≥1.5 or ≤–1.5; FDR ≤0.001). (C) HCA showing the top 12 DEGs in the *BEST1* c.851 A>G line compared to corrected 1 and 2. (D) Volcano plot illustration of DEGs between *BEST1 c.851A>G* vs corrected 1and 2. Genes likely important for macular RPE function and those implicated in Best disease pathology are highlighted and labeled. (E-F) Volcano plot depicting DEGs belonging to the TMEM and SLC gene families which are critical in ion transport, metabolite exchange, and cellular homeostasis in the RPE. A similar differential gene expression pattern was observed between *BEST1* c.851 A>G and corrected clones. The x-axis represents the log2 fold change expression between experimental groups, and the y-axis indicates the -log10 false discovery rate (FDR) value (adjusted p < 0.001). Genes upregulated in *BEST1* c.851 A>G RPE cells are shown in red, downregulated in blue, and non-significant in grey. Dashed lines denote thresholds for statistical significance and biologically relevant fold change (log2FC ≥ or ≤ 1.5).

Next, *BEST1* mutant RPE and corrected RPE monolayers were characterized for RPE-specific markers by immunofluorescence (IF) microscopy. In four independent differentiation experiments, the tight junction marker ZO-1 showed a similar localization pattern in both *BEST1* mutant RPE and corrected RPE cells (Figures 4A and 4B). Similarly, the apicobasal polarity marker EZRIN showed similar distribution in both *BEST1* mutant and corrected RPE cells (Figure 4A). In addition, PAX6, a transcription factor required for RPE development and melanogenesis, was expressed in RPE cells from both the *BEST1* mutant and corrected lines, (Supplementary Figure S5). Finally, BEST1, which is localized at the basolateral as well as the perinuclear region ^31^ displayed a punctate localization pattern in both *BEST1* mutant and corrected RPE cells (Figure 4B). However, BEST1 protein expression appeared reduced in mutant RPE cells compared to corrected RPE cells. Indeed, at 8 weeks of culture, western blot (WB) analysis revealed 8–11-fold reduced BEST1 protein levels in *BEST1* mutant compared to corrected RPE cells (p < 0.001) (Figure 5A). In addition, we observed two bands in the WB for BEST1 protein, which may represent alternative isoforms of BEST1. RPE65 protein was also decreased in the mutant RPE cells, with the corrected RPE cells showing approximately 4-5 fold greater protein levels (p < 0.005) (Figure 5B). Whereas Na⁺/K⁺-ATPase showed no difference (Figure 5C). Consistent with WB data, IF at 12 weeks of RPE culture revealed that mutant RPE cells displayed lower BEST1 and RPE65 protein levels compared with *BEST1* corrected RPE cells (Figure 5D).

**Figure 4:**
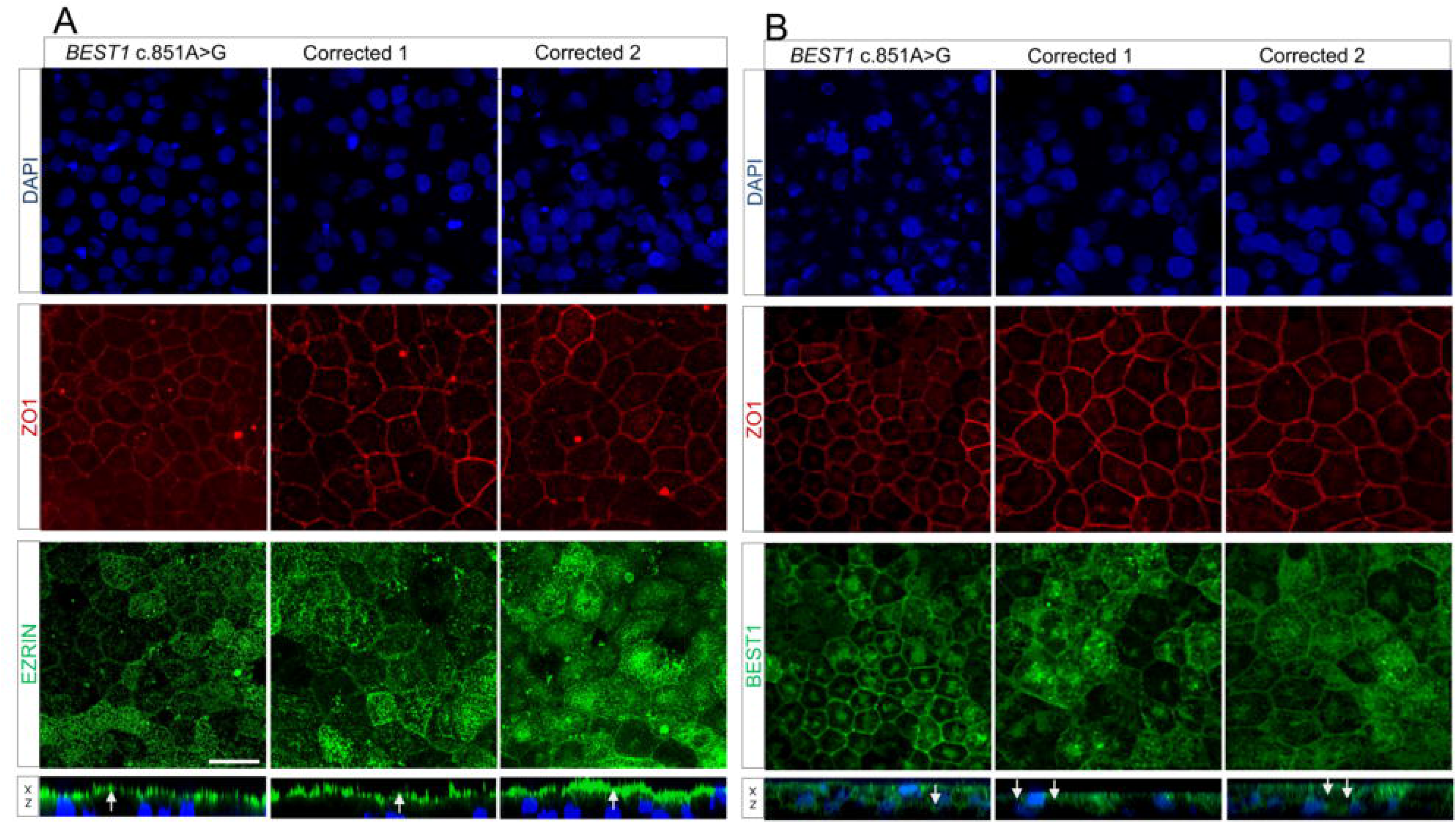
Characterization of RPE derived from *BEST1 c.851A>G* iPSCs and corrected clones. (A) Immunofluorescence staining (IF) shows expression of the apical polarity marker Ezrin (EZR) in RPE cells at week 4 of culture (post passage 1), enabling consistent comparison of marker localization between groups. XZ plane images demonstrate apical localization of EZR (white arrows) in the RPE cells of both genotypes. (B) Bestrophin-1 (BEST1) is similarly localized in cells of both genotypes. XZ plane images shows basolateral localization of BEST1 in RPE cells (white arrows), indicating expected cell polarity. A tight junctions marker, ZO1 was used to select morphologically comparable RPE cells for analysis. Representative image shown from 4 independent IF experiments. IF was performed in four independent RPE differentiation experiments. Scale bar = 20 µm.

**Figure 5:**
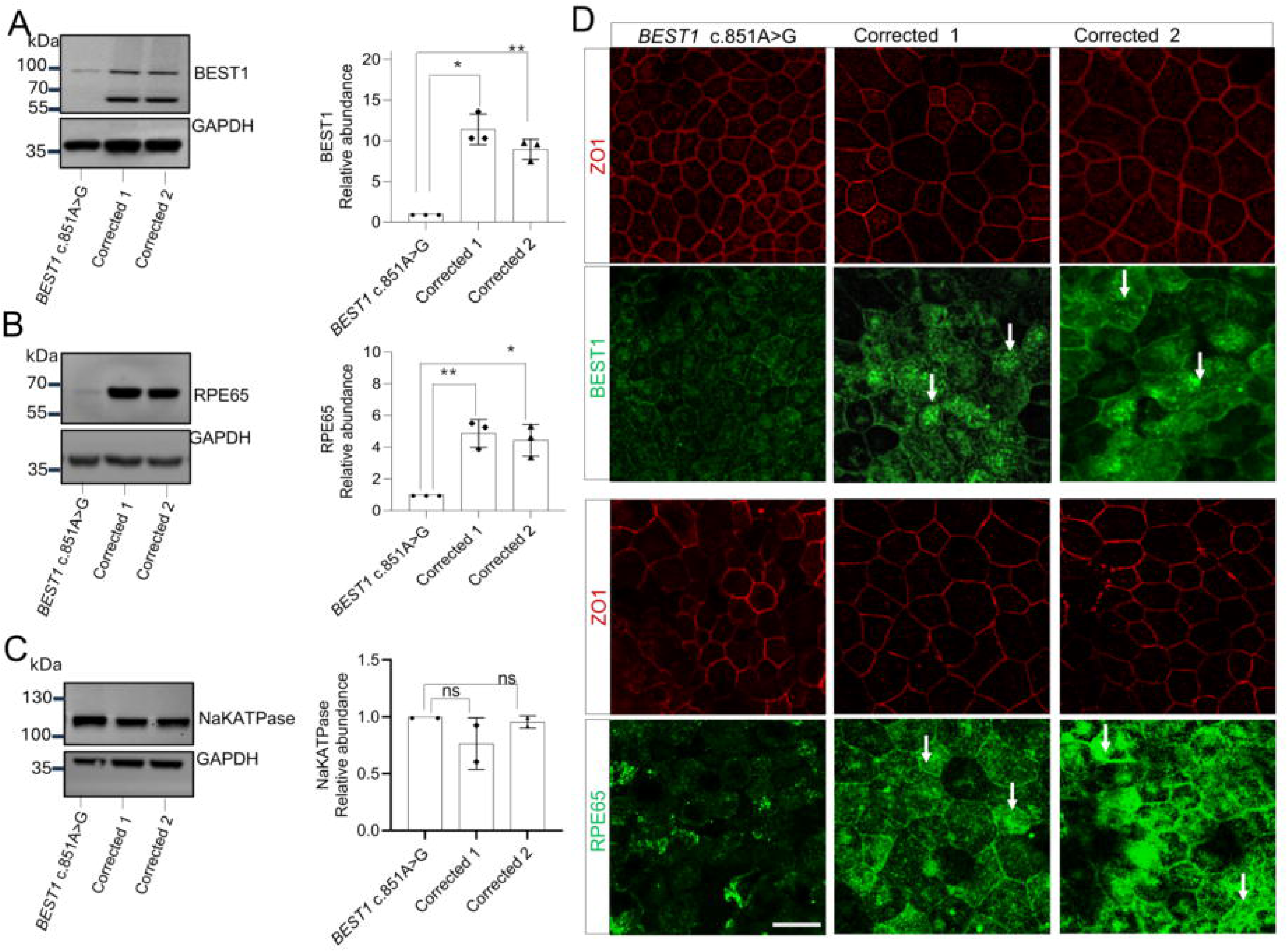
Comparative analysis of protein levels in RPE derived from *BEST1* c.851A>G iPSCs and corrected clones. Immunoblot at 8 weeks and quantification of band densitometry shows (A) reduced BEST1 protein levels in the *BEST1* mutant RPE cell line compared to corrected clones, (B) reduced RPE65 protein levels in the *BEST1* mutant RPE cell line compared to corrected clones, (C) sodium-potassium ATPase (Na⁺/K⁺-ATPase) protein levels is unaltered in the RPE derived from *BEST1* c. 851 A>G variant compared to corrected clones. Densitometry quantification of RPE65 and both bands for BEST1 (upper and lower band) indicate significant differences. The WB images shown are representative of the quantified data (Data represent n = 2–3 independent replicates from independent experiments). Protein extracts are from independent RPE differentiation experiments. Data are presented as mean ± SD; Statistically significant difference *p<0.05 and ** p<0.01. (D) Immunofluorescence microscopy to visualize BEST1 and RPE65 at 12 weeks. *BEST1* c. 851 A>G cells exhibit lower levels of BEST1 and RPE65 proteins compared to corrected clones, which show the expected punctate protein distribution (white arrows). ZO1 is used to select morphologically comparable regions Scale bar = 20 µm. Representative images shown (n=4). IF was performed in 2 independent RPE differentiation experiments

Transepithelial electrical resistance (TER), a measure of epithelial barrier integrity resulting from tight junction formation, and phagocytosis are key functional characteristics of RPE cells. We therefore assessed whether the *BEST1* c.851A>G variant impairs these functions. To evaluate TER, RPE cells were differentiated for 11 weeks, then passaged and cultured for an additional 9 weeks. TER measurements were measured at different time points starting from week 2 (Figure 6A and Supplementary Figure S6). At 2 weeks, no significant difference was observed between *BEST1* mutant RPE and corrected clones. Starting at 4 weeks, *BEST1* mutant RPE exhibited significantly lower TER compared to corrected RPE at all five time points, with an average difference ranging from ∼50 to 80 Ω·cm². We observed variability in TER between the corrected clones despite being derived from a single *BEST1* iPSC line (Supplementary Figure S6), suggesting that factors such as clonal heterogeneity or subtle culture related influences may contribute to variability. Overall, however, TER is higher in both corrected RPE compared to *BEST1* mutant RPE. These findings corroborate previous studies demonstrating that *BEST1* variants alter TER and thereby compromise the barrier properties of the RPE cells.^32,33^

**Figure 6:**
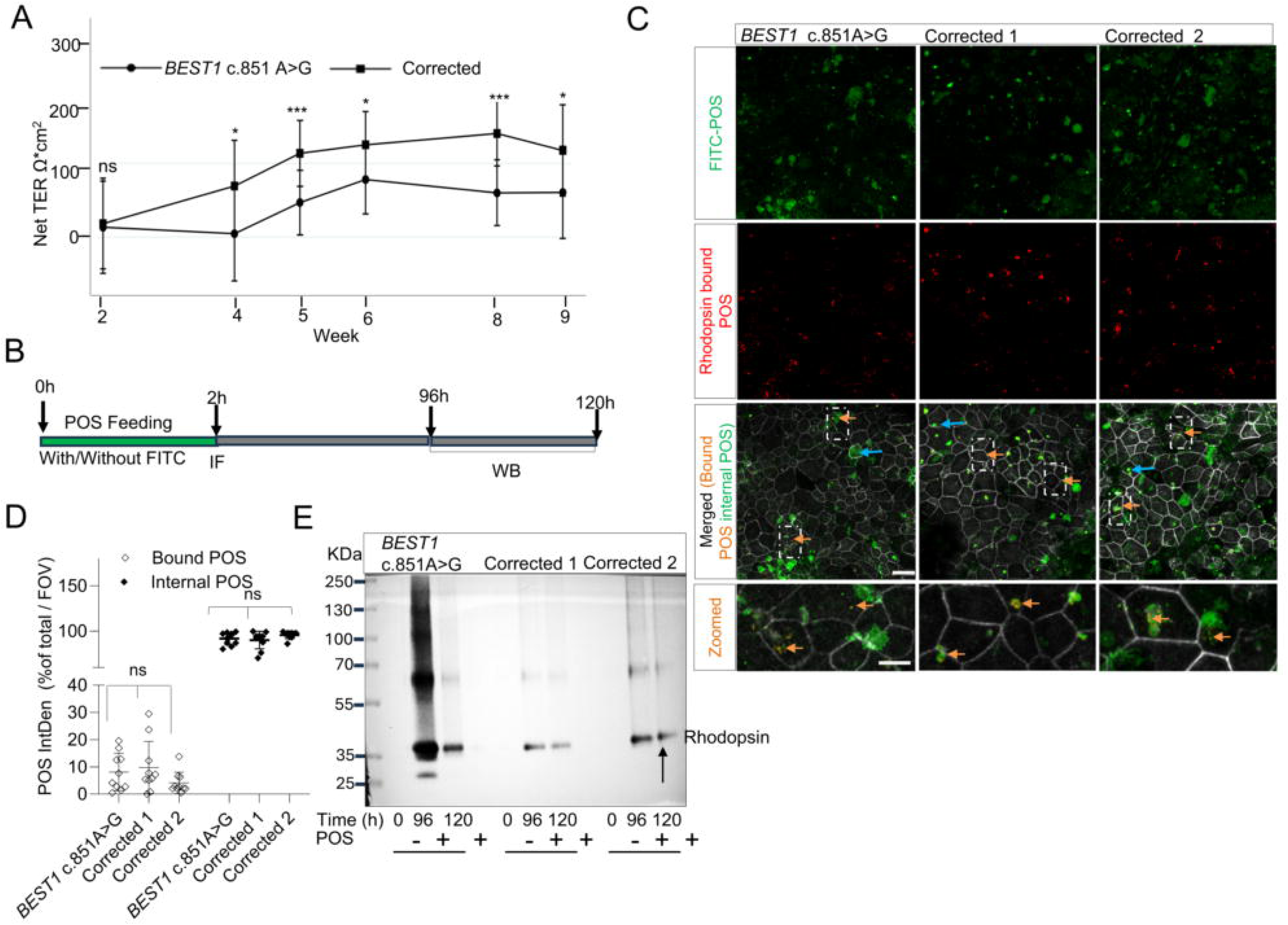
Functional rescue in RPE derived from the *BEST1* c.851A>G variant and corrected clones. (A) Transepithelial electrical resistance (TER) measurements of RPE cells showed higher TER in corrected clones compared to BEST1 mutant cells. At week 2, TER differences were not significant (p = 0.82). From week 4 onward, significant differences were detected (p = 0.02), largest at weeks 5 and 8 (p < 0.0001), and were still significant at weeks 6 and 9 (p = 0.02). Error bars represent SEM.* Represents a significant difference, and ns = non-significant (B) Schematic representation of evaluation timepoints for RPE mediated phagocytosis of photoreceptor outer segments (POS). (C-D) Representative confocal fluorescence images and quantification of POS uptake. POS were labeled with FITC (green). Surface-bound (external) POS were identified by labeling with anti-rhodopsin antibody B630 followed by Alexa Fluor 568. FITC fluorescence detected after exclusion of surface-bound signal (Rhodopsin bound POS) represents internal POS within the same field of view (indicated by blue arrow). Boxed regions zoom to show the rhodopsin bound POS (indicated by orange arrows) within the same field of view. Scale bar = 20 µm. (D) Quantification of POS fluorescence intensity was performed using Fiji (n=4; 2-3 ROI/sample, 2 independent experiments). Integrated density values were measured and expressed as a percentage of total POS integrated density per field of view (FOV). Data are presented as mean ± SD. ns = non-significant. (E) Western blot (WB) analysis of rhodopsin degradation in RPE cells. The results demonstrate a delay in rhodopsin degradation in *BEST1* c. 851A>G RPE compared with corrected RPE cells. The experiment was performed independently twice (n=4/ group), and a representative blot is shown.

To evaluate the phagocytic function of *BEST1* mutant and corrected RPE cells, we quantified the ability of the cells to bind, engulf, and degrade photoreceptor outer segments (POS) (Figure 6B). At 8 weeks of culture, *BEST1* mutant RPE and corrected RPE cells were challenged with FITC--labelled purified bovine rod POS for 2 hours before observing bound and internalized POS (Figure 6C). Quantification of POS revealed no significant difference between *BEST1* mutant RPE and corrected RPE cells in their ability to bind and internalize POS (Figure 6 D). We then evaluated the efficiency of POS clearance following phagocytic binding and engulfment at 8 weeks of culture. As rhodopsin constitutes approximately 80–90% of the total protein content in rod POS, we used WB to assess rhodopsin degradation as a readout of degradation efficiency. Notably, we observed far more rapid rhodopsin degradation in the corrected RPE compared to *BEST1* mutant RPE cells at 96 and 120 h post POS feeding, indicated by WB band intensity (Figure 6E, representative of two independent experiments (n=4/ group). Taken together, our findings suggest that although the rate of POS engulfment is unaffected in *BEST1* mutant RPE, the efficiency of POS degradation is reduced.

### Correction of the BEST1 c.851A>G variant in RPE monolayers

Given the functional differences observed in mutant versus corrected RPE cells, we evaluated whether the pathogenic *BEST1* c.851A>G variant could be efficiently and selectively corrected in RPE monolayers. *BEST1* mutant RPE monolayers were co-transfected with plasmids expressing the sgRNA and CBE4-SpG (P2A–EGFP) using Lipofectamine (Figure 7A). Sanger sequencing confirmed editing at the *BEST1* locus (Figures 7B and 7C), with bulk sequencing revealed the highest editing at c.851 (55.5 ± 10.6%) with lower editing at the bystander c.853 (34.9 ± 23.2%). The resulting bystander c.853 (coding strand G>A substitution (p. Val285Ile), represents a chemically conservative change at a residue that is poorly conserved across species. Computational predictions classified the induced bystander Val285Ile variant as benign by PolyPhen-2 and tolerated by SIFT (score 0.39) (Supplementary Figures S7A–E). These results suggest that CBE-mediated base editing can edit the *BEST1* c. 851A>G variant in mutant RPE monolayers and that any bystander edits are unlikely to alter BEST1 function.

**Figure 7:**
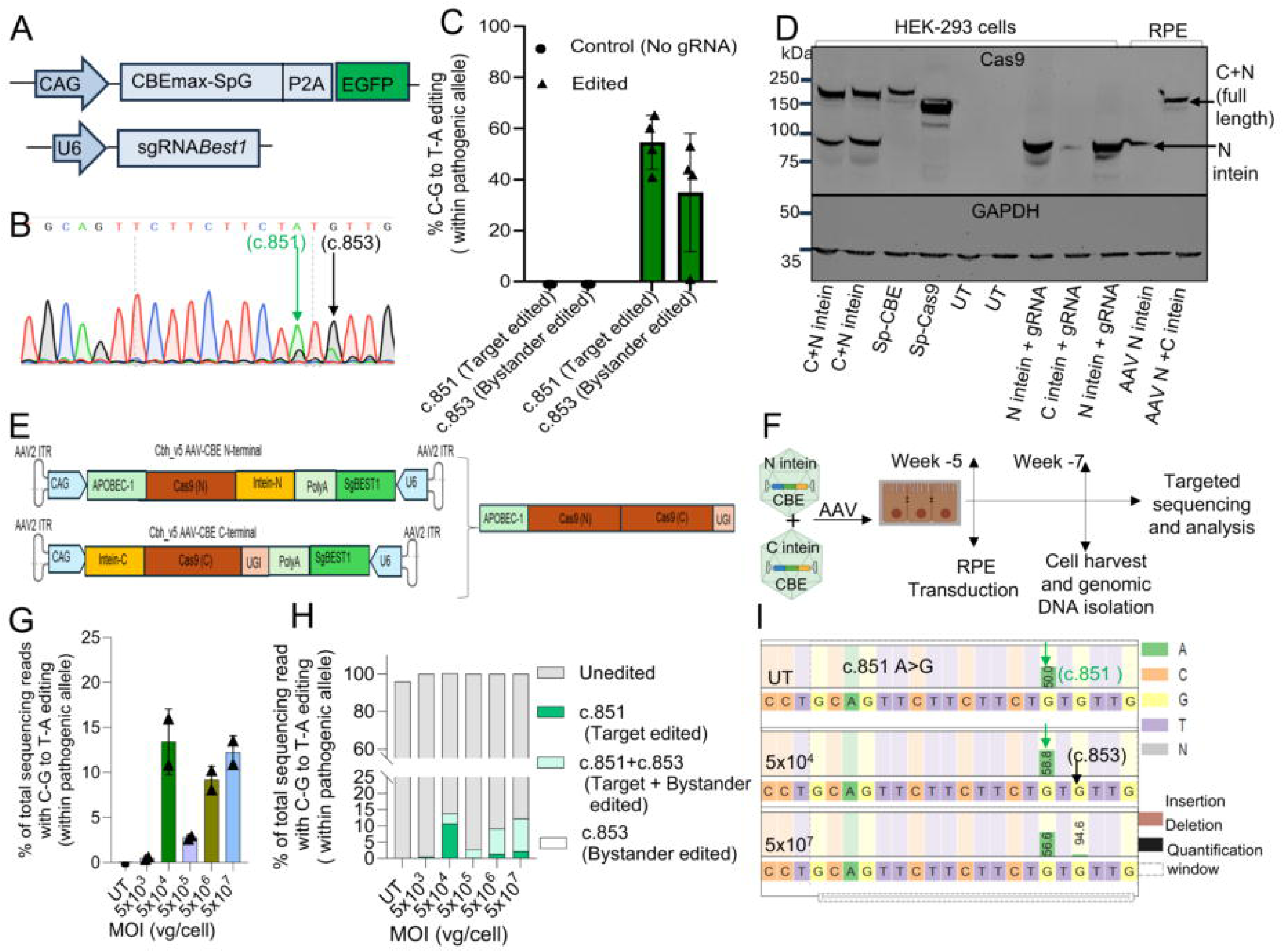
*BEST1* (c. 851 A>G) editing in RPE cells. (A) Schematic of CBE4max and sgRNA expression cassettes. (B) Sanger sequencing confirms the presence of the edited *BEST1* allele; correct edit (c.851, green arrow) and bystander edit (c.853, black arrow) demonstrating correction of the pathogenic allele. (C) Quantification of editing *BEST1* efficiency in Lipofectamine-transfected RPE cells (n=4). (D) Western blot showing abundance of the full-length CBE protein in HEK293 transfected cells with full-length CBE, Cas9 and intein (CBE) expressing plasmid and in RPE cells transduced with AAV split-inteins. (E) Schematic of the AAV intein reconstitution strategy. (F) Schematic of RPE transduction and time frame of the analysis. (G) On-target (c.851) editing efficiency at each AAV intein (C and N) dose. The edited nucleotide (C-T) confirms AAV-mediated base correction of *BEST1* c.851A>G allele. (H) Summary graph showing base editing efficiencies at positions c.851 and c.853 across varying AAV doses. Both cytosines (c.851 and c.853) were quantified to assess AAV dose dependent editing changes within the target window. Allele frequencies below 1% of total reads were excluded from the calculation of C-G-to-T-A editing efficiency. (I) Quantification of nucleotide frequencies within the editing window at MOI 5x10^4^ (dose with highest editing efficiency) and at MOI 5x10^7^ (highest tested dose). The sgRNA target position is marked by a gray dashed line. The targeted nucleotide “G” (indicated black) upon editing changes to “A” (indicated green) are shown for the coding (+) strand spanning the *BEST1* c.851 A>G. MOI = multiplicity of infection (vector genomes (vg) /cell).

### BEST1 editing via dual AAV split-intein

Because the packaging capacity of AAV (∼4.5 kb) is insufficient to deliver the full-length base editor, we utilized C- and N-terminal inteins in two AAVs (Figures 7E and 7F) to deliver split CBE-Cas9 as previously described.^34^ First, we confirmed the expression of full-length CBE after split-intein plasmid transfection in HEK293 cells followed by dual AAV transduced mutant RPE cells (Figure 7D). Mutant RPE cells transduced with both AAVs (1:1 ratio) at doses ranging from multiplicity of infection (MOI, vg/cell) 5x10^3^ to 10^7^ (Figure 7F), were analyzed for C-to-T base editing at: 1) target base only (c.851); 2) both target base and bystander base (c.851 and c.853); 3) bystander base only (c.853); and 4) unedited. The total editing efficiency for c.851 (intended site) ranged from 0.5 ± 0.1% to 13.4 ± 3.6%, with the highest editing efficiency (13.4 ± 3.6%) achieved at MOI 5x10^4^ (Figure 7G). The editing efficiencies for the target base only (c.851) ranged from 0.5 ± 0.1% to 10.6 ± 8.4%, with best results also achieved at MOI 5x10^4^ (Figure 7H). Bystander only (c.853) editing was not observed at any of the tested AAV doses (Figure 7I). These findings demonstrate correction of the pathogenic allele using CBE delivery via dual AAV.

## DISCUSSSION

In this study, we established a personalized base-editing strategy for BVMD. Our findings show that patient-specific iPSC-derived RPE cells effectively model Best disease associated with the *BEST1* c.851A>G variant, demonstrated by increased cell death, transcriptomic changes, reduced RPE-specific protein levels, reduced barrier integrity, and inefficient POS degradation. These phenotypes were normalized in isogenic corrected lines, underscoring the pathogenic role of this variant and its amenability to gene correction. We then demonstrated successful editing of mutant RPE monolayers using a split-intein approach.

The *BEST1* c.851A>G (p. Tyr284Cys) variant disrupts RPE survival and function (Figures 2-6). Reports suggest that BVMD may partly result from reduced levels of functional BEST1 protein^8,35,36,37,21^ and we did observe reduced BEST1 protein (Figure 5) consistent with tyrosine 284 lying within a conserved C-terminal region important for channel stability. Its substitution with a smaller, reactive cysteine would likely disrupt folding and trafficking and consequently reduce protein stability and function. This dysfunction may then cause secondary transcriptional changes that also affect the key RPE pathways.^38^ Several dysregulated genes uncovered here overlap with those previously implicated in RPE maturation, phagocytosis, and metabolic regulation (Figure 3E and 3F). This convergence suggests that *BEST1* dysfunction may drive broader effects on RPE function beyond its canonical role in chloride channel activity. Further investigation of molecular pathway perturbations and the mechanistic basis of cell death caused by this and other *BEST1* variants could uncover therapeutic targets for restoring RPE function in patients with BVMD.

This study has identified important areas for future investigation and certain limitations. First, the immunoblotting (Figure 4A) indicates two bands of BEST1 protein, which we believe are BEST1 isoforms or alternatively, a higher molecular weight oligomer of BEST1. Importantly, these observations do not rule out the possibility that BEST1 exists physiologically as a dimer and that the minimal functional unit is a dimer.^39^ However, we did not examine BEST1 isoforms in the current study, and this remains an interesting question for future investigation. Second, the reduced abundance of RPE fate proteins in the mutant may result from the pathological effects of the mutant BEST1 protein on RPE development and maturation.^40,35^ BEST1 mutant RPE showed reduced protein and mRNA levels compared to corrected RPE cells, suggesting effects on both RNA abundance and protein stability (Figures 4 and 5). Third, transcriptomic profiling was done at a single time point (week 8), limiting insight into dynamic changes over time. Future investigation should incorporate longitudinal sampling and single cell RNA sequencing to resolve cellular heterogeneity and better understand the restoration within corrected RPE populations.^44^

Finally, there remain issues with editing efficiency and specificity that could impact the translational potential of this approach. Delivery of the full-length CBE via lipofection achieved ≥50% editing efficiency after sorting for GFP+ cells, whereas the dual AAV-based split-intein CBE strategy resulted in maximum editing efficiency of 13% in patient-derived RPE cells (Figure 7). This editing efficiency is consistent with prior reports, but it would likely need to be higher to achieve clinical efficacy^41,42,43^. Still, the proportion of cell rescue required to alter disease trajectory for patients with BVMD and the c.851A>G variant specifically remains unknown.

While our dual AAV-based delivery system shows early promise, further optimization of delivery will be critical to enhance editing outcomes and achieve greater therapeutic effects. In addition, the proposed base editing strategy has not yet been tested across multiple genetic backgrounds. Future studies should assess robustness in a diverse set of patient-derived iPSC lines to validate the applicability of this cytidine base editing approach in the *BEST1* gene. Moreover, eliminating bystander edits may be necessary for clinical translation, as even low-frequency modifications affecting poorly conserved amino acids (Supplementary Figures S7A– E) could have unpredictable consequences in RPE cells. This underscores the need for rigorous biochemical and functional characterization of bystander edited cells to validate that editing does not affect RPE function.

In conclusion, our findings support the translational potential of base editing as a therapeutic strategy for dominant retinal diseases, particularly when combined with scalable and clinically relevant AAV-based delivery platforms. These results lay the groundwork for future *in vivo* validation and highlight the broader applicability of base editing to correct pathogenic variants in dominant IRDs.^44–46^ Moreover, corrected RPE offers a potential source for autologous cell therapy in advanced stages of Best disease caused by various *BEST1* variants.

## MATERIALS AND METHODS

### Study approval and human subjects

The ocular disease-focused exome sequencing was performed at Children’s Hospital Los Angeles (CHLA) at the Center of Personalized Medicine as part of routine clinical care. Local sequence information from the exome data was obtained under a research protocol approved by the CHLA Institutional Review Board (CHLA-18-00467). Patient blood was obtained, and the genotype determined by Sanger sequencing confirmed the *BEST1* c.851A>G variant previously identified in the clinical test. All experimental protocols for iPSC use and generation from patient blood were approved under CHLA IRB (IRB-CHLA-19-00369).

### iPSC generation

Erythrocyte progenitor cells were isolated from the participant’s peripheral blood using SepMate^TM^ tubes (catalog no. 86415, Stem Cell Technologies, Vancouver, BC, Canada) and the RosetteSep™ Human Progenitor Cell Basic Pre-Enrichment Kit (catalog no. 05924, Stem Cell Technologies, catalog no. 05924, Stem Cell Technologies) as per instructions. To enrich for erythroid progenitors, cells were cultured in StemSpan^TM^ SFEM II media (catalog no.09605, Stem Cell Technologies, atalog no. 05924, Stem Cell Technologies) containing StemSpan^TM^ Erythroid Expansion Supplement (SSEE) (catalog no.02692, Stem Cell Technologies, Vancouver, BC, Canada) for nine days. Flow cytometry using anti-CD235a (catalog no.559943, BD Biosciences, San Jose, CA, USA) and anti-CD71 (catalog no.555537, BD Biosciences, San Jose, CA, USA) was used to verify erythroid progenitor enrichment. For reprogramming, the erythroid progenitors were transfected with episomal plasmids expressing SOX2, KLF4, LMYC, Lin28, OCT3/4, shp53 and EBNA (catalog no.27078, 27080, 27077, 27624, Addgene, Watertown, MA, USA) using the Neon^TM^ Transfection System and cultured on Matrigel (catalog no. BD354277, Corning, NY, USA) in SSEE media for 2 days. On days 3-7 media was transitioned to ReproTeSR^TM^ medium (catalog no. 05926, Stem Cell Technologies, Vancouver, BC, Canada). After 18-27 days, resultant colonies exhibiting pluripotent stem cell morphology were manually isolated and cultured on Matrigel in mTeSR^TM^ Plus (catalog no. 100-1130, Stem Cell Technologies, Corning, NY, USA)). Subsequently, the iPSC clones were passaged either manually or using ReLeSR^TM^ (catalog no. 05872, Stem Cell Technologies, Vancouver, BC, Canada). iPSC between passages 5-8 were tested for the following quality control tests: 1) Stemness marker and pluripotency, 2) Recurrent copy number variations (CNV) (iCS-digital™ PSC 24-probe test, Stem Genomics, Montpellier, France), 3) and karyotype (G-banding) at CHLA Cytogenetics lab.

### Characterization of iPSC pluripotency and stemness

The undifferentiated state of iPSCs was determined using antibodies to the surface marker SSEA4 (APC anti-human SSEA4 antibody, catalog. no. 330418, BioLegend, San Diego, CA, USA) and the transcription factor OCT3/4 (PE anti-human/mouse OCT3/4 antibody, REAfinity™, catalog. no. 130-120-236, Miltenyi Biotec, Bergisch Gladbach, Germany). Staining was performed using the transcription factor staining buffer set (catalog. no. 130-122-981, Miltenyi Biotec, Bergisch Gladbach, Germany) according to the manufacturer’s protocol with minor modifications. The fixation/permeabilization solution was prepared by mixing solution 1 and solution 2 at a 1:4 ratio, and the permeabilization buffer was diluted 1:10 with deionized water. Cells were washed 1–2 times with 1× PBS and incubated with Accutase (catalog no. A6964, STEMCELL Technologies, Vancouver, BC, Canada) for 10 min at 37°C. After detachment, cells were washed, resuspended in 1× PBS at a density of 1 × 10⁶ cells/mL, and centrifuged at 300 × g for 5 min. The cell pellet was resuspended in 3 mL of cold fixation/permeabilization buffer and incubated for 30 min at 37°C. Subsequently, 5 mL of cold PBS was added, and cells were centrifuged at 300 × g for 5 min at 4°C. The pellet was washed with 1 mL of permeabilization buffer and centrifuged again at 300 × g for 5 min. Cells were then resuspended in 96 µL of permeabilization buffer per 1 × 10⁶ cells and divided into two aliquots (48 µL each). 4′,6-diamidino-2-phenylindole (DAPI,5 µg/mL in 1× PBS) only used as a control, and the other was stained with 1 µL of OCT3/4 antibody and 2.5 µL of APC antihuman SSEA4 antibody. Following incubation on ice for 30 min, cells were washed with 1 mL of permeabilization buffer and centrifuged at 300 × g for 5 min. The final pellet was resuspended in 300 µL of 5% FBS in PBS and analyzed on a BD LSR II flow cytometer.

Following FACS analysis, iPSC lines were evaluated for the expression of pluripotency markers OCT4, SSEA4, NANOG, and TRA-1-81 (catalog no. 9656S, Cell Signaling, Danvers, MA, USA) by immunofluorescence. Cells were cultured in µ-Slide 8 Well ibiTreat chambers (ibidi, catalog. No. 80826) to approximately 70% confluency. Cells were rinsed twice with 400 µL of 1× phosphate-buffered saline (PBS) and fixed with 200 µL of 4% paraformaldehyde (PFA) for 15 min at room temperature (RT). After fixation, cells were washed and incubated with 200 µL of antibody Incubation Solution (AIS) containing 3% horse serum and 0.3% Triton X-100 in 1× PBS (prepared using (Diethyl pyrocarbonate (DEPC)-treated water) for 1 h at RT Primary antibodies were diluted 1:200 in Blocking Buffer (5% normal horse serum in 1× PBS prepared with DEPC-treated water). Cells were incubated with 200 µL of primary antibody solution overnight at 4°C, followed by 1 h at RT. Wells were rinsed twice with 1× PBS and incubated with Alexa fluor conjugated secondary antibodies (1:500 in blocking buffer) for 1 h at RT. Nuclei were counterstained with 200 µL of DAPI for 10 min at RT. Samples were rinsed twice with PBS and imaged using a fluorescence microscope.

### Plasmid vector construction and AAV production

Single-guide RNAs (sgRNAs) were designed to target the *BEST1* c.851A>G variant using CHOPCHOP (https://choppchop.cbu.uib.no) and CRISPOR (https://crispor.gi.ucsc.edu) and CRISPick (https://portals.broadinstitute.org/gppx/crispick/public) tools. Sense and antisense oligonucleotides used for sgRNA (Supplementary Table 1) were synthesized by Integrated DNA Technologies (IDT, Coralville, IA, USA). Oligos were annealed and ligated into the BbsI (catalog no. R0539S, NEB, Ipswich, MA, USA) cut site of the digested pSPgRNA plasmid (catalog no.47108, Addgene, Watertown, MA, USA) using T4 DNA ligase (catalog no. M0202S, NEB, Ipswich, MA, USA)) and plasmid DNA generated by transforming NEB 5 alpha competent E. coli (catalog no.C2987I, NEB, Ipswich, MA, USA)) with the ligated product. The expression plasmid of the cytosine base editor, pCAG-CBE4max-SpG-P2A-EGFP (RTW4552) (catalog no.139998, Addgene, Watertown, MA, USA), was used to create C-T transitions. For AAV vector construction, *BEST1* sgRNA was first PCR amplified from the previously generated pSPgRNA-BEST1 vector, and inserted into NotI (catalog no.R3189S, NEB, Ipswich,MA,USA) and XhoI (catalog no. R0146S, NEB, Ipswich,MA,USA)) double digested Cbh_v5 AAV-CBE N-terminal as well as C-terminal plasmids (catalog no.137175 and catalog no137176 Addgene, Watertown, MA,USA) using Gibson assembly (catalog no. GA1100-10 NEB, Ipswich, MA, USA). All constructs were verified by whole plasmid sequencing at Plasmid Saurus (https://Plasmidsaurus.com) and the sequences are provided in supplementary information (plasmid sequence 1). Primers used for cloning are listed in supplementary Table 2. AAV vectors were packaged into the AAV2.7m8 capsid (catalog no. 64839, Addgene, Watertown, MA,USA)), and AAV was produced using standard triple transfection followed by purification and titer quantification using primers and probe set (Supplementary Table) as previously described. The yields of N-intein and C-intein vectors were 2.06 × 10¹³ vg/mL and 3.11 × 10¹³ vg/mL, respectively.

### Isogenic iPSC line generation

iPSCs were cultured at 60-70% confluence in mTeSR^TM^ Plus culture media. Single cell suspensions were prepared using Accutase (catalog no.07920, STEMCELL Technologies, Vancouver, BC, Canada). Final cell suspension of 2 x 10^5^cells/ 10 ul was prepared in buffer R (catalog no. MPK1096, Invitrogen, Carlsbad, CA, USA). CBE (pCAG-CBE4max-SpG-P2A-EGFP) and sgRNA-expressing plasmid at a molar ratio of 1:1 (total 3 ug of DNA in 1-2 ul) was added to the cell suspension. Cells were transfected using a Neon Transfector (cat. No MPK5000, Thermo Fisher Scientific, Waltham, MA, USA) with the following parameters: voltage 1400 V, pulse width 20 ms, and pulse number 1. Transfected cells were cultured in 24-well plates in mTeSR™ medium supplemented with Celone-R2 (catalog no. 100-0691, STEMCELL Technologies, Vancouver, BC, Canada). 48 hours post-transfection, cells were FACS-sorted, and genomic DNA was isolated using the NucleoSpin (catalog no. 740100.50, Takara Bio, Kusatsu, Shiga, Japan). The *BEST1* c.851A>G locus was amplified by PCR using primers listed in Supplementary Table 4 and the PCR Master Mix (catalog no. RR330A, Takara Bio, Kusatsu, Shiga, Japan). PCR amplicons were purified using the NucleoSpin Gel and PCR Clean-up Kit (catalog no. 740609.5, Takara Bio, Kusatsu, Shiga, Japan), and genotyping was performed by Sanger sequencing (GENEWIZ). After confirming base editing in the cell population (Supplementary Figure S2), GFP positive single cells were isolated using FACS, cultured, expanded, and genotyped. Correctly edited clones were characterized for pluripotency, karyotype (CHLA cytogenetic facility), and single copy number variation (SNV) (Stem Genomics, Inc.), (Supplementary Figure S1)

### Off-target base editing analysis

The top ten potential off-target sites for the *BEST1* sgRNA were identified by an *in-silico* tool using IDT CRISPR-Cas9 gRNA checker (CRISPR-Cas9 guide RNA design checker | IDT (idtdna.com)). The parameters used were an NGG/PAM, and up to four mismatches to the sgRNA sequence (Supplementary Table 4). Genomic DNA was isolated from the *BEST1* mutant and corrected iPSC lines using NucleoSpin DNA RapidLyse (catalog no.740100, Takara Kusatsu, Shiga, Japan), followed by PCR using primers specific to off-target sites for site amplification and Sanger sequencing. For Sanger sequencing, each amplicon was gel purified using NucleoSpin® Gel and PCR Clean-Up kit (catalog no.740609, Takara, Kusatsu, Shiga, Japan) eluted in 10 µl ultrapure water (Millipore) and quantified by NanoDrop (catalog no 13400518, Thermo Fisher Scientific, Waltham, MA, USA)). The Sanger sequencing premix was prepared by adding 1.5 µl eluted DNA (∼ 40 ng) and 2.5 µl 10 µM sequencing primers. Off-target primers are listed in Supplementary Table 5.

### Directed differentiation of iPSC to RPE

*BEST1* c.851A>G and corrected isogenic iPSC lines were differentiated into RPE following a previously published protocol with modifications.^26^ Briefly, iPSCs were differentiated over 11 weeks, and cells were passaged at weeks 2, 7, and 11. Nicotinamide (NIC; catalog no. 72340, Sigma-Aldrich, St. Louis, MO, USA) was added at a final concentration of 10 mM for 14 days following each passage, and cells were treated with deoxyribonuclease I (DNase I; catalog no. D4527, Sigma-Aldrich, St. Louis, MO, USA)) at each passage.

### Transepithelial resistance (TER) measurements

At passage 1 (week11), RPE cells were plated in transwell inserts (catalog no. 07-200-154, Corning, Corning Incorporate, Corning, NY, USA) at a density of 1x10^5^ cells per 0.33 cm². Transepithelial resistance (TER) of RPE monolayer cultured on transwell was measured using EVOM2 (catalog no. EVOM2, World Precision Instruments) according to the manufacturer’s instructions and following published protocol.^47^ TER was calculated by subtracting the blank measurement obtained from cell-free matrigel-coated filter and multiplying the difference by the area of the transwell.

### BEST1 base editing in RPE cells

To edit *BEST1* by transfection, 2x10^5^ RPE cells were plated on a matrigel coated 24-well plate (catalog no. 10861-700, VWR, Randoe, PA, USA) before transfection. Cells were transfected, after 48h with CBE (catalog no.139998, Addgene, Watertown, MA,USA) base editor plasmid sgRNA expression plasmid (catalog no. 47108, Addgene, Watertown, MA,USA) at a molar ratio of 1:1 (total 3 ug) using Lipofectamine 3000 (catalog no. L3000001, Life Technologies, Carlsbad, CA,USA) prepared in Opti-MEM I (catalog no. 31985062, Fisher Scientific,Waltham, MA,USA). 24h post-transfection media were replaced, and at 48h post-transfection EGFP expressing cells were FACS sorted. Genomic DNA was isolated, PCR amplified, and subjected to Sanger sequencing as described above. Base editing efficiency was quantified with EditR (https://moriaritylab.shinyapps.io/editr_v10/).^48^

To edit *BEST1* by AAV-mediated transduction of CBE inteins (C and N intein), RPE cells were plated in 48-well plates at 2 × 10⁵ cells/well on matrigel coated surfaces. At week 5 of culture, cells were transduced with AAV at doses ranging from 1x10^9^-1x10^12^ vg. 24h post-transduction, the culture medium was replaced, and cells were maintained for an additional 2 weeks. At the end of 7 weeks, DNA was isolated as described above (*iPSC line generation*).

### Targeted amplicon sequencing analysis

For high-throughput sequencing (HTS), the genomic region flanking each targeted *BEST1* locus was PCR-amplified using primers listed in Supplementary Table 6, with partial Illumina adapters provided by Amplicon-EZ added to the 5’ end of each forward and reverse primer. A unique 6–8 nucleotide barcode was incorporated into the PCR product via adapter sequences added to the primers, enabling discrimination of amplicons from different repeats or experimental conditions pooled within the same sample.

PCR was performed in 10 µl reactions with 50 pmol of each primer, 10 ∼ 50 ng template, and 5 µl 2× Phanta Max master mix (Dye Plus, Vazyme). The annealing temperature was set to 60 °C, and 35 cycles were used for amplification. The desired DNA fragment was extracted from the PCR mix using buffer PB (catalog no 19066, Qiagen,Hilden, Germany) when a single and clear band was expected or extracted using QIAquick PCR & Gel Cleanup Kit (catalog no 28506, Qiagen, Hilden,Germany) after Tris-acetate EDTA (catalog no.T9650, Sigma Aldrich) agarose gel electrophoresis. The FASTQ files from NGS amplicon sequencing were analyzed by CRISPResso2 (http://crispresso.pinellolab.partners.org/).

### RNA sequence analysis

RNA was extracted from RPE cells that had been cultured on 24-well transwell inserts (1x10^5^/well) for 8 weeks by RNeasy Kits (catalog no ID:74104, Qiagen, Hilden, Germany). Total RNA was quantified by Qubit and integrity assessed on a TapeStation. Azenta Life Sciences (South Plainfield, NJ, USA) performed RNA library preparation and sequencing as follows: RNA-seq libraries were prepared with the NEBNext Ultra II RNA Library Prep Kit (Illumina) per manufacturer’s instructions. Poly(A) mRNA was enriched, fragmented, and reverse-transcribed to double-stranded cDNA. Libraries were end-repaired, A-tailed, adapter-ligated, indexed, and PCR-amplified. Library quality and concentration were evaluated by TapeStation, Qubit, and qPCR. Prepared libraries were multiplexed and sequenced on an Illumina NovaSeq with 2×150 bp paired-end reads. Base calling was performed with NovaSeq Control Software; BCL files were converted to FASTQ and demultiplexed with bcl2fastq, allowing one index mismatch. RNA-seq data was analyzed using Partek™ Flow™ software, v11.0. Briefly, raw sequencing reads were trimmed based on quality scores (Phred score >= 35, minimum trimmed read length = 25nt) and mapped to GRCh38 using STAR (2.7.8a, Dobin *et al.* 2013) with Gencode v42 as a guide. The aligned reads were then quantified to Gencode v42 using the Partek E/M method. Genes with less than 10 reads in any samples were excluded from the downstream analysis. DESeq2 (Love MI, Huber W, Anders S; 2014) was used for differential gene expression analysis and differentially expressed genes (DEGs) were selected using cutoffs FDR<0.001 and fold changes greater than 1.5 in either direction.

### Western blot analysis

To assess expression of Cas9-CBE intein proteins from expression vectors, HEK293T cells (catalog no. 632180, Takara,Kusatsu, Shiga,Japan) were cultured in 24-well plates at a density of 4 × 10⁴ cells per well. 24h after seeding, cells were transfected with the plasmids pCAG-CBE4max-SpG-P2A-EGFP, Cbh_v5 AAV-CBE N, and Cbh_v5 AAV-CBE C using Lipofectamine, following the manufacturer’s instructions. 48h post-transfection, cells were harvested using trypsin, and whole-cell lysates were prepared as below. To assess Cas9-CBE intein expression in RPE cells transduced with AAV, *BEST1* mutant RPE cells were cultured in 48 well plates (2 × 10⁵ cells/well) for 4 weeks, transduced, and incubated for an additional 2 weeks before harvesting. Similarly, *BEST1* mutant and corrected RPE cells were cultured on transwell inserts for 8 weeks before harvest.

For protein extraction, cells were lysed in ice cold RIPA buffer (catalog no. 89900, Thermo Fisher, Waltham,Ma,USA) supplemented with protease inhibitor (catalog no. 5892970001, Sigma, St.Louis, MO, USA) and phosphatase inhibitor (catalog no. 4906845001, Sigma, St.Louis, MO, USA). Lysates were incubated on ice for 30 minutes and centrifuged at 14,000 rpm for 20 minutes at 4°C. The resulting supernatant was collected for WB analysis. Total protein concentration was quantified using the Qubit Protein Quantification Kit (catalog no. Q33211, Thermo Fisher,Waltham, MA,USA). The protein lysate was mixed with 6X loading dye and held at 95°C for 5 minutes. 25 μg of protein per lane were separated using a NuPAGE 4-12% Bis-Tris gel (catalog no.NP0322BOX, Invitrogen, Carlsbad, CA, USA) and transferred onto PVDF membrane (catalog no.GE10600023, Sigma Aldrich, St. Louis, MO, USA)), followed by 1h blocking in 5% nonfat milk (catalog no.1706404, Bio-Rad, Hercules, CA, USA) in tris-buffered saline (TBS) (catalog no. J62938.K7, Thermo Fisher, Waltham, MA, USA) containing 0.075% Tween 20 (TBS-T). The membrane was incubated with primary antibody diluted in 3% nonfat milk in TBS-T overnight at 4°C. Membranes were washed 3 x 10min with TBS-T and then incubated with anti-mouse IgG-HRP antibody for 1 h at room temperature. After 3 x 10min wash with TBS-T chemiluminescent Substrate (catalog no.32106, ThermoFisher, Waltham, MA, USA) was added and membrane visualized using iBright 1500 (Invitrogen, Carlsbad, CA, USA).

### Immunofluorescent staining (IF)

RPE cells cultured on transwells were rinsed twice with 200 µL of 1X PBS and fixed with 200 µL 4% paraformaldehyde (PFA) for 15 minutes at room temperature (RT). Following fixation, cells were washed and incubated with 200 µL of AIS (3% horse serum, 0.3% Triton X-100 in 1X PBS prepared with DEPC water) in a humidified chamber for 1 hour at RT. After a subsequent wash with 1X PBS, 100 µL of primary antibody in AIS was added to the corresponding wells and incubated at 4°C for 16h on a rotator. After incubation, the wells were rinsed with 1X PBS three times and incubated with AlexaFluor-conjugated secondary antibodies in AIS for 1h at RT. Cells were then stained with 5 µg/mL of DAPI, prepared in 1X PBS, for 15min at RT. The well linings were carefully removed from the transwell, and the cut section mounted using ProLong™ Diamond Antifade Mountant (catalog no. P36970 Fisher Scientific, Waltham, MA, USA) on a glass slide, and the slide sealed. Primary and secondary antibodies used in IF and WB are listed in the supplementary Table 7.

### Confocal fluorescence microscopy and quantification

Confocal Z-stack images were acquired using an inverted STELLARIS 5 laser scanning confocal system equipped with a white-light laser and 63×/1.40 HC PL APO CS2 oil immersion objective lens (Leica Microsystems). Fluorophores were excited with laser lines at 405 nm (DAPI), 488 nm (Alexa Fluor 488), 555 nm (Alexa Fluor 568), and 639 nm (Alexa Fluor 647). Images were collected at a resolution of 512 × 512 pixels per frame, corresponding to a field of view of either 92.39 × 92.39 µm² or 184.58 × 184.58 µm², depending on acquisition settings. Z-stacks were acquired with a step size of 1 µm, and all imaging parameters were kept constant within each experiment. Laser power and detector gain were adjusted to optimize dynamic range while avoiding signal saturation. The pinhole diameter was calculated for 1 Airy unit at 580 nm and held constants for all channels. For quantitative analysis of POS fluorescence intensity, identical regions of interest (ROIs) were applied across all images and samples, and maximum intensity projections of optical sections acquired at 63× magnification were analyzed using FIJI (ImageJ, NIH).

### Apoptosis assay

RPE cells were plated on the transwell (1x10^5^ cells / well) and cultured for 4 or 8 weeks. Apoptosis assay using the Annexin-V FITC/PT kit (catalog no.88.8005-72, Life Technology, Carlsbad, CA, USA) according to the manufacturer’s instructions. Briefly, cells were incubated with 200 μL of 1 × binding buffer containing 5 μL of Annexin V-FITC and 5 μL of PI 15 min in the dark at RT then analyzed by flow cytometry (BD FACSCalibur, NJ, USA). Annexin V+ /PI− cells were considered early apoptotic cells, annexin V+ /PI+ cells were considered late apoptotic cells, and annexin V-/PI+ were necrotic. The apoptotic cell population comprises the sum of early and late apoptosis.

### Phagocytosis assays

RPE cells were cultured on transwell inserts for 8 weeks, and two different assays, IF and WB, were performed to evaluate phagocytosis and photoreceptor outer segment (POS) degradation function of the RPE cells, respectively. In the IF assay, RPE cells were fed FITC-labeled POS 10 pos/cell (InVision Bioresources, Sanford, ME, USA,) supplemented with 2µg/mL protein S (catalog no. HCPS 0090 Hyphen Biomed, Seoul, South Korea)) and 2.5µg/mL MFGE8 (catalog no.2767-MF Bio-Techne, Minneapolis, MN, USA) for 2 h. Cells were washed five times with PBS (Ca+, Mg+) (catalog no. 14040133, Gibco,Waltham, MA, USA) by vigorous pipetting to remove any unbound POS and IF was performed to check the bound and internal POS using anti-Rhodopsin antibody, following published protocol.^49,50^

POS degradation was evaluated by WB analysis. Briefly, RPE cells (plated at 10^6^ cells / well) were fed unlabeled POS for 2 h, followed by thorough washing to remove unbound POS. Unfed RPE cultures were used as controls for all experiments. RPE cells were then maintained in culture for an additional 96 or 120 h. At each time point, cells were harvested, and total protein lysates were prepared to assess POS degradation by RPE cells. WB was performed using unboiled samples to preserve rhodopsin integrity and allow detection of degradation. 25 μg of total protein quantified using Qubit Protein Assay Kits (catalog no. Q33211, WA, Massachusetts, USA) was loaded per lane, following the WB protocol described above.

### Statistical analysis

Adjusted mean TER values from *BEST1 c.851A>G* and two corrected clones were compared at various weeks of growth using a linear mixed effects (LME) model with restricted maximum likelihood estimation and a degrees of freedom correction.^51^ Each RPE differentiation was treated as a random effect to account for batch differences; data from two corrected clones were combined in one group for analysis. Western blot band intensities were quantified by densitometry using ImageJ and normalized to the loading control, and statistical significance was determined using a two-tailed Student’s t-test. For phagocytosis and apoptosis assays, statistical significance was assessed using a t-test with Benjamini–Krieger and Yekutieli multiple comparison corrections. Data in the text and figures are presented as mean ± SD.

## Data availability statement

RNA-seq data have been deposited in the NCBI Gene Expression Omnibus (GEO) under accession number GSE300866.

## Acknowledgements

The authors are grateful to the patient who participated in this study. We also acknowledge the core facilities at The Saban Research Institute of Children’s Hospital Los Angeles, including the Flow Cytometry (FACS) Core for cell-sorting assistance, the Imaging Core for providing key instrumentation and technical support, and the Stem Cell Core for iPSC generation and characterization.

## Author contributions

**SK**: Conceptualization, Investigation, Methodology, Analysis, Writing original draft, Funding acquisition, Review & Editing, and Project management

**JGA**: Conceptualization, Investigation, Methodology, Review, Editing, and Project management

**KS, AB, MH, PG**: Methodology, Review & editing

**AS, NH:** iPSC generation and characterization

**JB, MR**: Review & Editing

**GEF:** Images analysis, Editing, and Review

**ML**: Bioinformatics, Editing, and Review

**RJS**: Investigation, conceptualization, computational prediction of variant, and strategy Review & Editing.

**CA:** Writing, Review & Editing

**DC**: Funding acquisition, Review & Editing.

**AN:** Conceptualization, Funding acquisition, Project management, Review & Editing.

## Declaration of Interests

A.N. serves as a consultant for AAVantgarde, Alia Tx, Astellas, Opus Genetics, Beacon, Biogen, Blackstone, Blue Gen Therapeutics Foundation, BlueRock Therapeutics, Cellio, Epicrispr, Eyebiotech, Immuneering, Novartis, Sunregen, Ultragenyx, and Vida Ventures; received grants or contracts from Atsena, Johnson & Johnson, Spark, Opus Genetics, BlueRock, and AAVantgarde; patent pending for Lats Kinase Inhibitor to Treat Retinal Degeneration; holds stocks or stock options in Cellio and Raresight. The other authors have no interests to declare.

## Notes

This research was supported in part by: an unrestricted grant to the Department of Ophthalmology at the Keck School of Medicine of USC from Research to Prevent Blindness, New York, NY, USA (A.N., C.A., D.C.), Las Madrinas Endowment in Experimental Therapeutics for Ophthalmology (A.N.), the Knight Templar Eye Foundation (A.N., C.A., D.C.), and the CIRM Training Program for Stem Cell and Regenerative Medicine Research (EDUC4-12802) at Children’s Hospital Los Angeles (S.K.).

