## Supplementary material for "Allele-specific correction of dominant Best vitelliform macular dystrophy in patient-derived retinal pigment epithelium": Supp Data

**Supplementary Table 1–** gRNA oligos sequence

| | |
|---|---|
| F (BEST1)- gRNA1 | Phos 5'-CACCGCAACACAGAAGAAGAACTGC-3' |
| R (BEST1)-gRNA1 | Phos 5'- AAACGCAGTTCTTCTTCTGTGTTGC--3' |
| F (BEST1)-gRNA2 | Phos 5'-CACCAACACAGAAGAAGAACTGA-3' |
| R (BEST1)-gRNA2 | Phos 5'-AAACTCAGTTCTTCTTCTGTGTT-3' |

**Supplementary Table 2-** Primer cloning (Gibson assembly)

| **Gene name** | **BEST1** |
|---|---|
| Primer name/ seqF 5'-3' | CCAACTCCATCACTAGGGGTTCCTGCGAGGGCCTATTTCCCATGATTC |
| Primer name/seq R 5'-3' | CTGGGGATGCGGTGGGCTCTATGGCAAAAAAAGCACCGACTCGGTG |

**Supplementary Table 3-** Probe and primer for AAV titer quantification

| | |
|---|---|
| Probe 5'-3' | 56-FAM/CAAGTGTAT/ZEN/CATATGCCAAGTACGCCCC/3IABkFQ |
| Primer1 5'-3' | GTCCCATAAGGTCATGTACTGG |
| Primer2 5'-3' | TCAATGGGTGGAGTATTTACGG |

**Supplementary Table 4-** Identified off-target (OFT) for *BEST1*sgRNA

| | | | | | |
|---|---|---|---|---|---|
| CAACACAGAAGAAGAACTGC | AGG | 0 | 0 | BEST1 | chr11:-155799529- |
| CACC-CAGAAGAAGAACTGC | TGG | 4 | 2 | EPS8 | chr12:-15704212 |
| CAAC-CA-AAGAAGAACTGC | TAG | 8 | 2 | KIAA1549L | chr11:+33399352 |
| CACCACA-AAAAAGAACTGC | AGG | 8 | 3 | FOXJ3 | chr1:+42245360 |
| CATC-CAGGAGAAGAACTGC | TGG | 10 | 3 | TRAF3 | chr14:-102883087 |
| CAGCTCACAAGAAGGACTGC | AAG | 11 | 4 | Intergenic | chr3:+53960687 |
| CAGGACAGTAAGAAGAACTGC | TGG | 13 | 3 | SUGT1P4 | chr9:-97240805 |

| | | | | | |
|---|---|---|---|---|---|
| TAGCACATAAAAAGAACTGC | TAG | 13 | 4 | LINC02498 (long intergenic non-protein coding RNA 2498) | chr4:-10747098 |
| CACCCACAGATGAAGAACTGC | TAG | 15 | 3 | OBI1-AS1 | chr13:+78311240 |
| TCACACAGAAGCAGAACTGC | TGG | 17 | 3 | XKR9 | chr8:+70804163 |
| GGACACAGAGAAAGAACTGC | AGG | 17 | 4 | ZNF462 | chr9:-106890483 |

**Supplementary Table 5-** Primer for off-target (OFT) analysis

| OFT | Primer name/ seqF 5'-3' | Primer name/seq R 5'-3' | amplicon size |
|---|---|---|---|
| EPS8 | GCACGCATGCCAAGGAAAT | AATCTGCCCACCACTTACCT | 950 |
| KIAA1549L | AGGGGAACGCTTTTCTCCTC | GCTGGGGGATTCAGAACCAT | 322 |
| FOXJ3 | CTTGGGCATGGTGTATTA | ACACTGACAGCCACACTAATC | 853 |
| TRAF3 | ACCCTATGGGAAAGCATACGTG | TTGAAAAGCACAGGCCGGG | 900/901 |
| Intergenic | TCTGCCAACTACAGCACATGA | CTCTCCAAAGCACCTACCCTCG | 591 |
| SUGT1P4 | TATAGGTGTGACCATGCCCG | TGCCACGTCATGCTGTTACT | 642 |
| LINC02498 | TCTCCCAGAGCACCCTTTTC | TCCAGCCCTCAAGGTGTTTG | 624 |
| OBI1-AS1 | ACATCCTGCAGGTCACCAGT | CCTGTGCAATAGCAGGGAAC | 723 |
| XKR9 | GCAGGTATATCCTGGGACATTAG | GATCATCCCACTTTGCACATAAA | 363 |
| ZNF462 | GAGTGCAGGGTAACTGCCTT | GAGCATATGCCCTCTGACCC | 730 |

**Supplementary Table 6 -**Primer for the *BEST1* locus (High throughput sequencing (HTS)). Partial Illumina adapters sequence (lower case) added at the 5' end of the primer.

| Gene name | Primer name/ seq F 5'-3' | Primer name/seq R 5'-3' |
|---|---|---|
| BEST1 (HTS) | **acactctttccctacacgacgctcttccgatct** CCCCTGGAGCATCCTGATTT | **gactggagttcagacgtgtgctcttccgatct** CTGGCTGCTTTCTACCCGTG |
| BEST1 | CAACACAGAAGAAGAACTGC | CCCCTGGAGCATCCTGATTT |

**Supplementary Table 7** -Antibodies and conditions used for immunofluorescence staining (IF) and Western blot (WB)

| **Antibody** | **Source** | **Catalogue no** | **Dilution** | **IF/WB** |
|---|---|---|---|---|
| ZO1 | Invitrogen | 40-2200 | 1:300 | IF |
| ZO1 | Invitrogen | 339100 | 1:300 | IF |
| Ezrin | Atlas | HPA021616 | 1 100 | IF |
| BEST1 | Abcam | ab2182 | 1:300 | IF |
| PAX6 | BioLegend | 901301 | 1:100 | IF |
| RPE65 | Abcam | ab235950 | 1:150 | IF |
| anti Rabbit AF488 | Invitrogen, | A21206 | 1:1000 | IF |
| anti Mouse AF488 | Invitrogen, | A21202 | 1:1000 | IF |
| anti Rabbit AF568 | Invitrogen, | A10042 | 1:1000 | IF |
| anti MouseAF568 | Invitrogen, | A10037 | 1:1000 | IF |
| BEST1 | Abcam | ab22182 | 1:1000 | WB |
| RPE65 | Abcam | ab13826 | 1-1000 | WB |
| NaKATP | Santacrutz | sc-58628 | 1-1000 | WB |
| GAPDH | Cell signaling | 5174S | 1:1,000 | WB |
| Beta-actin | Cell signaling | 4967S | 1:1,000 | WB |
| Cas9 | ThermoFisher | MA5-23519 | 1:1000 | WB |
| Anti-mouse,IgG-HRP | Cell signaling | 7076P2 | 1:10,000 | WB |
| Anti-mouse,IgG-HRP | Cell signaling | 7074P2 | 1:10,000 | WB |

**Plasmid sequence**

**1** – pSP-gRNA – sequence highlighted in yellow

1. sgRNA1- Sanger sequence plasmid (catalog no.47108, Addgene)

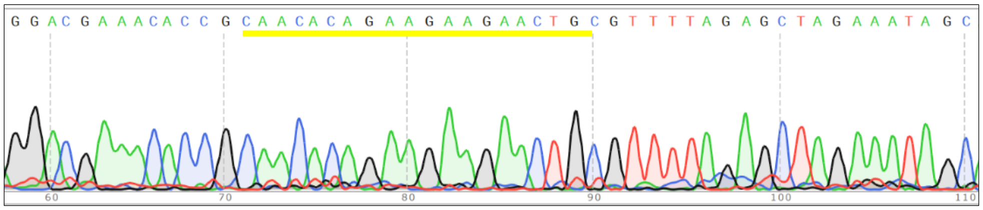


2. sgRNA2-Sanger sequence plasmid (catalog no.47108, Addgene)

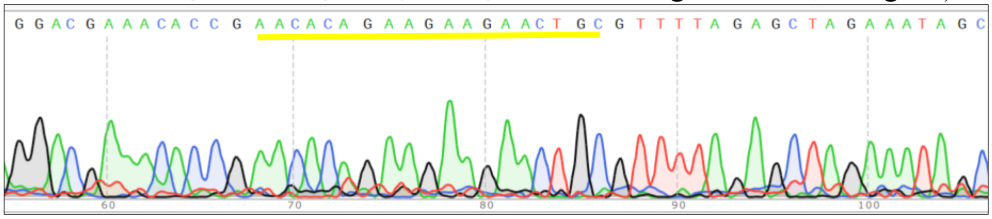


**Plasmid sequence**

**1) SpAAV-N intein (sgRNA sequence)**

TTGAGATCCTTTTTTTCTGCGCGTAATCTGCTGCTTGCAAACAAAAAAACCACCGCTA
CCAGCGGTGGTTTGTTTGCCGGATCAAGAGCTACCAACTCTTTTTCCGAAGGTAACTG
GCTTCAGCAGAGCGCAGATACCAAATACTGTTCTTCTAGTGTAGCCGTAGTTAGGCCA
CCACTTCAAGAACTCTGTAGCACCGCCTACATACCTCGCTCTGCTAATCCTGTTACCA
GTGGCTGCTGCCAGTGGCGATAAGTCGTGTCTTACCGGGTTGGACTCAAGACGATAG
TTACCGGATAAGGCGCAGCGGTCGGGCTGAACGGGGGGTTCGTGCACACAGCCCAG
CTTGGAGCGAACGACCTACACCGAACTGAGATACCTACAGCGTGAGCTATGAGAAAG
CGCCACGCTTCCCGAAGGGAGAAAGGCGGACAGGTATCCGGTAAGCGGCAGGGTCG
GAACAGGAGAGCGCACGAGGGAGCTTCCAGGGGGAAACGCCTGGTATCTTTATAGTC
CTGTCGGGTTTCGCCACCTCTGACTTGAGCGTCGATTTTTGTGATGCTCGTCAGGGGG
GCGGAGCCTATGGAAAAACGCCAGCAACGCGGCCTTTTTACGGTTCCTGGCCTTTTG
CTGGCCTTTTGCTCACATGTTCTTTCCTGCGTTATCCCCTGATTCTGTGGATAACCGTAT
TACCGCCTTTGAGTGAGCTGATACCGCTCGCCGCAGCCGAACGACCGAGCGCAGCGA
GTCAGTGAGCGAGGAAGCGGAAGAGCGCCCAATACGCAAACCGCCTCTCCCCGCGC
GTTGGCCGATTCATTAATGCAGCTGGCACGACAGGTTTCCCGACTGGAAAGCGGGCA
GTGAGCGCAACGCAATTAATGTGAGTTAGCTCACTCATTAGGCACCCCAGGCTTTACA

CTTTATGCTTCCGGCTCGTATGTTGTGTGGAATTGTGAGCGGATAACAATTTCACACAG
GAAACAGCTATGACCATGATTACGCCAGATTTAATTAAGGCTGCGCGCTCGCTCGCTC
ACTGAGGCCGCCCGGGCAAAGCCCGGGCGTCGGGCGACCTTTGGTCGCCCGGCCTC
AGTGAGCGAGCGAGCGCGCAGAGAGGGAGTGGCCAACTCCATCACTAGGGGTTCCT
GCGGCCTCTAGATCAGGGTACCCGTTACATAACTTACGGTAAATGGCCCGCCTGGCTG
ACCGCCCAACGACCCCCGCCCATTGACGTCAATAGTAACGCCAATAGGGACTTTCCAT
TGACGTCAATGGGTGGAGTATTTACGGTAAACTGCCCACTTGGCAGTACATCAAGTGT
ATCATATGCCAAGTACGCCCCCTATTGACGTCAATGACGGTAAATGGCCCGCCTGGCA
TTGTGCCCAGTACATGACCTTATGGGACTTTCCTACTTGGCAGTACATCTACGTATTAG
TCATCGCTATTACCATGGTCGAGGTGAGCCCCACGTTCTGCTTCACTCTCCCCATCTCC
CCCCCCTCCCCACCCCCAATTTTGTATTTATTTATTTTTTAATTATTTTGTGCAGCGATG
GGGGCGGGGGGGGGGGGGGGCGCGCGCCAGGCGGGGCGGGGCGGGGCGAGGGG
CGGGGCGGGGCGAGGCGGAGAGGTGCGGCGGCAGCCAATCAGAGCGGCGCGCTCC
GAAAGTTTCCTTTTATGGCGAGGCGGCGGCGGCGGCGGCCCTATAAAAAGCGAAGCG
CGCGGCGGGCGGGAGTCGCTGCGCGCTGCCTTCGCCCCGTGCCCCGCTCCGCCGCCG
CCTCGCGCCGCCCGCCCCGGCTCTGACTGACCGCGTTACTCCCACAGGTGAGCGGGC
GGGACGGCCCTTCTCCTCCGGGCTGTAATTAGCTGAGCAAGAGGTAAGGGTTTAAGG
GATGGTTGGTTGGTGGGGTATTAATGTTTAATTACCTGGAGCACCTGCCTGAAATCAC
TTTTTTTCAGGTTGGACCGGTGCCACCATGAAACGGACAGCCGACGGAAGCGAGTTC
GAGTCACCAAAGAAGAAGCGGAAAGTCTCCTCAGAGACTGGGCCTGTCGCCGTCGA
TCCAACCCTGCGCCGCCGGATTGAACCTCACGAGTTTGAAGTGTTCTTTGACCCCCG
GGAGCTGAGAAAGGAGACATGCCTGCTGTACGAGATCAACTGGGGAGGCAGGCACT
CCATCTGGAGGCACACCTCTCAGAACACAAATAAGCACGTGGAGGTGAACTTCATCG
AGAAGTTTACCACAGAGCGGTACTTCTGCCCCAATACCAGATGTAGCATCACATGGTT
TCTGAGCTGGTCCCCTTGCGGAGAGTGTAGCAGGGCCATCACCGAGTTCCTGTCCAG
ATATCCACACGTGACACTGTTTATCTACATCGCCAGGCTGTATCACCACGCAGACCCA
AGGAATAGGCAGGGCCTGCGCGATCTGATCAGCTCCGGCGTGACCATCCAGATCATG
ACAGAGCAGGAGTCCGGCTACTGCTGGCGGAACTTCGTGAATTATTCTCCTAGCAAC
GAGGCCCACTGGCCTAGGTACCCACACCTGTGGGTGCGCCTGTACGTGCTGGAGCTG
TATTGCATCATCCTGGGCCTGCCCCCTTGTCTGAATATCCTGCGGAGAAAGCAGCCCC
AGCTGACCTTCTTTACAATCGCCCTGCAGTCTTGTCACTATCAGAGGCTGCCACCCCA

CATCCTGTGGGCCACAGGCCTGAAGTCTGGAGGATCTAGCGGAGGATCCTCTGGCAG
CGAGACACCAGGAACAAGCGAGTCAGCAACACCAGAGAGCAGTGGCGGCAGCAGC
GGCGGCAGCGACAAGAAGTACAGCATCGGCCTGGCCATCGGCACCAACTCTGTGGG
CTGGGCCGTGATCACCGACGAGTACAAGGTGCCCAGCAAGAAATTCAAGGTGCTGG
GCAACACCGACCGGCACAGCATCAAGAAGAACCTGATCGGAGCCCTGCTGTTCGAC
AGCGGCGAAACAGCCGAGGCCACCCGGCTGAAGAGAACCGCCAGAAGAAGATACA
CCAGACGGAAGAACCGGATCTGCTATCTGCAAGAGATCTTCAGCAACGAGATGGCCA
AGGTGGACGACAGCTTCTTCCACAGACTGGAAGAGTCCTTCCTGGTGGAAGAGGATA
AGAAGCACGAGCGGCACCCCATCTTCGGCAACATCGTGGACGAGGTGGCCTACCAC
GAGAAGTACCCCACCATCTACCACCTGAGAAAGAAACTGGTGGACAGCACCGACAA
GGCCGACCTGCGGCTGATCTATCTGGCCCTGGCCCACATGATCAAGTTCCGGGGCCAC
TTCCTGATCGAGGGCGACCTGAACCCCGACAACAGCGACGTGGACAAGCTGTTCATC
CAGCTGGTGCAGACCTACAACCAGCTGTTCGAGGAAAACCCCATCAACGCCAGCGG
CGTGGACGCCAAGGCCATCCTGTCTGCCAGACTGAGCAAGAGCAGACGGCTGGAAA
ATCTGATCGCCCAGCTGCCCGGCGAGAAGAAGAATGGCCTGTTCGGAAACCTGATTG
CCCTGAGCCTGGGCCTGACCCCCAACTTCAAGAGCAACTTCGACCTGGCCGAGGATG
CCAAACTGCAGCTGAGCAAGGACACCTACGACGACGACCTGGACAACCTGCTGGCC
CAGATCGGCGACCAGTACGCCGACCTGTTTCTGGCCGCCAAGAACCTGTCCGACGCC
ATCCTGCTGAGCGACATCCTGAGAGTGAACACCGAGATCACCAAGGCCCCCCTGAGC
GCCTCTATGATCAAGAGATACGACGAGCACCACCAGGACCTGACCCTGCTGAAAGCT
CTCGTGCGGCAGCAGCTGCCTGAGAAGTACAAAGAGATTTTCTTCGACCAGAGCAAG
AACGGCTACGCCGGCTACATTGACGGCGGAGCCAGCCAGGAAGAGTTCTACAAGTTC
ATCAAGCCCATCCTGGAAAAGATGGACGGCACCGAGGAACTGCTCGTGAAGCTGAA
CAGAGAGGACCTGCTGCGGAAGCAGCGGACCTTCGACAACGGCAGCATCCCCCACC
AGATCCACCTGGGAGAGCTGCACGCCATTCTGCGGCGGCAGGAAGATTTTTACCCAT
TCCTGAAGGACAACCGGGAAAAGATCGAGAAGATCCTGACCTTCCGCATCCCCTACT
ACGTGGGCCCTCTGGCCAGGGGAAACAGCAGATTCGCCTGGATGACCAGAAAGAGC
GAGGAAACCATCACCCCCTGGAACTTCGAGGAAGTGGTGGACAAGGGCGCTTCCGC
CCAGAGCTTCATCGAGCGGATGACCAACTTCGATAAGAACCTGCCCAACGAGAAGGT
GCTGCCCAAGCACAGCCTGCTGTACGAGTACTTCACCGTGTATAACGAGCTGACCAA
AGTGAAATACGTGACCGAGGGAATGAGAAAGCCCGCCTTCCTGAGCGGCGAGCAGA

AAAAGGCCATCGTGGACCTGCTGTTCAAGACCAACCGGAAAGTGACCGTGAAGCAG
CTGAAAGAGGACTACTTCAAGAAAATCGAGTGCCTGTCCTACGAGACAGAGATCCTG
ACAGTGGAGTATGGCCTGCTGCCAATCGGCAAGATCGTGGAGAAGAGGATCGAGTGT
ACCGTGTACTCTGTGGATAACAATGGCAACATCTATACACAGCCCGTGGCACAGTGGC
ACGATAGGGGAGAGCAGGAGGTGTTCGAGTATTGCCTGGAGGACGGCAGCCTGATC
AGGGCAACCAAGGACCACAAGTTCATGACAGTGGATGGCCAGATGCTGCCCATCGAC
GAGATTTTCGAGCGGGAGCTGGACCTGATGAGAGTGGATAACCTGCCTAATAGCGGA
GGCAGTAAAAGAACAGCAGACGGGAGTGAGTTTGAGCCCAAGAAAAAGAGAAAGG
TGTAAGATCTGATAATCAACCTCTGGATTACAAAATTTGTGAAAGATTGACTGGTATTC
TTAACTATGTTGCTCCTTTTACGCTATGTGGATACGCTGCTTTAATGCCTTTGTATCATG
CTATTGCTTCCCGTATGGCTTTCATTTTCTCCTCCTTGTATAAATCCTGGTTAGTTCTTG
CCACGGCGGAACTCATCGCCGCCTGCCTTGCCCGCTGCTGGACAGGGGCTCGGCTGT
TGGGCACTGACAATTCCGTGGTGCGACTGTGCCTTCTAGTTGCCAGCCATCTGTTGTT
TGCCCCTCCCCCGTGCCTTCCTTGACCCTGGAAGGTGCCACTCCCACTGTCCTTTCCT
AATAAAATGAGGAAATTGCATCGCATTGTCTGAGTAGGTGTCATTCTATTCTGGGGGG
TGGGGTGGGGCAGGACAGCAAGGGGGAGGATTGGGAAGACAATAGCAGGCATGCTG
GGGATGCGGTGGGCTCTATGGCAAAAAAAGCACCGACTCGGTGCCACTTTTTCAAGT
TGATAACGGACTAGCCTTATTTTAACTTGCTATTTCTAGCTCTAAAACGCAGTTCTTCT
TCTGTGTTGCGGTGTTTCGTCCTTTCCACAAGATATATAAAGCCAAGAAATCGAAATA
CTTTCAAGTTACGATAAGCATATGATAGTCCATTTTAAAACATAATTTTAAAACTGCAA
ACTACCCAAGAAATTATTACTTTCTACGTCACGTATTTTGTACTAATATCTTTGTGTTTA
CAGTCAAATTAATTCCAATTATCTCTCTAACAGCCTTGTATCGTATATGCAAATATGAAG
GAATCATGGGAAATAGGCCCTCGCAGGAACCCCTAGTGATGGAGTTGGCCACTCCCT
CTCTGCGCGCTCGCTCGCTCACTGAGGCCGGGCGACCAAAGGTCGCCCGACGCCCG
GGCTTTGCCCGGGCGGCCTCAGTGAGCGAGCGAGCGCGCAGCCTTAATTAACCTAAT
TCACTGGCCGTCGTTTTACAACGTCGTGACTGGGAAAACCCTGGCGTTACCCAACTT
AATCGCCTTGCAGCACATCCCCCTTTCGCCAGCTGGCGTAATAGCGAAGAGGCCCGC
ACCGATCGCCCTTCCCAACAGTTGCGCAGCCTGAATGGCGAATGGACGCGCCCTGT
AGCGGCGCATTAAGCGCGGCGGGTGTGGTGGTTACGCGCAGCGTGACCGCTACACTT
GCCAGCGCCCTAGCGCCCGCTCCTTTCGCTTTCTTCCCTTCCTTTCTCGCCACGTTCGC
CGGCTTTCCCCGTCAAGCTCTAAATCGGGGGCTCCCTTTAGGGTTCCGATTTAGTGCT

TTACGGCACCTCGACCCCAAAAAACTTGATTAGGGTGATGGTTCACGTAGTGGGCCAT
CGCCCTGATAGACGGTTTTTCGCCCTTTGACGTTGGAGTCCACGTTCTTTAATAGTGG
ACTCTTGTTCCAAACTGGAACAACACTCAACCCTATCTCGGTCTATTCTTTTGATTTAT
AAGGGATTTTGCCGATTTCGGCCTATTGGTTAAAAAATGAGCTGATTTAACAAAAATT
TAACGCGAATTTTAACAAAATATTAACGCTTACAATTTAGGTGGCACTTTTCGGGGAA
ATGTGCGCGGAACCCCTATTTGTTTATTTTTCTAAATACATTCAAATATGTATCCGCTCA
TGAGACAATAACCCTGATAAATGCTTCAATAATATTGAAAAAGGAAGAGTATGAGTAT
TCAACATTTCCGTGTCGCCCTTATTCCCTTTTTTGCGGCATTTTGCCTTCCTGTTTTTGC
TCACCCAGAAACGCTGGTGAAAGTAAAAGATGCTGAAGATCAGTTGGGTGCACGAG
TGGGTTACATCGAACTGGATCTCAACAGCGGTAAGATCCTTGAGAGTTTTCGCCCCGA
AGAACGTTTTCCAATGATGAGCACTTTTAAAGTTCTGCTATGTGGCGCGGTATTATCCC
GTATTGACGCCGGGCAAGAGCAACTCGGTCGCCGCATACACTATTCTCAGAATGACTT
GGTTGAGTACTCACCAGTCACAGAAAAGCATCTTACGGATGGCATGACAGTAAGAGA
ATTATGCAGTGCTGCCATAACCATGAGTGATAACACTGCGGCCAACTTACTTCTGACA
ACGATCGGAGGACCGAAGGAGCTAACCGCTTTTTTGCACAACATGGGGGATCATGTA
ACTCGCCTTGATCGTTGGGAACCGGAGCTGAATGAAGCCATACCAAACGACGAGCGT
GACACCACGATGCCTGTAGCAATGGCAACAACGTTGCGCAAACTATTAACTGGCGAA
CTACTTACTCTAGCTTCCCGGCAACAATTAATAGACTGGATGGAGGCGGATAAAGTTG
CAGGACCACTTCTGCGCTCGGCCCTTCCGGCTGGCTGGTTTATTGCTGATAAATCTGG
AGCCGGTGAGCGTGGGTCTCGCGGTATCATTGCAGCACTGGGGCCAGATGGTAAGCC
CTCCCGTATCGTAGTTATCTACACGACGGGGAGTCAGGCAACTATGGATGAACGAAAT
AGACAGATCGCTGAGATAGGTGCCTCACTGATTAAGCATTGGTAACTGTCAGACCAA
GTTTACTCATATATACTTTAGATTGATTTAAAACTTCATTTTTAATTTAAAAGGATCTAG
GTGAAGATCCTTTTTGATAATCTCATGACCAAAATCCCTTAACGTGAGTTTTCGTTCCA
CTGAGCGTCAGACCCCGTAGAAAAGATCAAAGGATCTTC

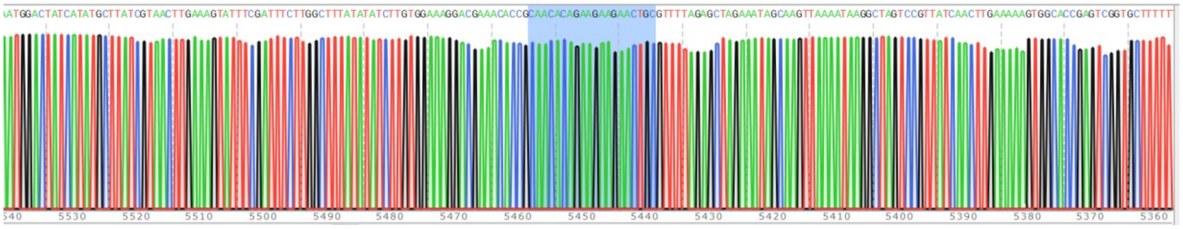

Sanger trace file – gRNA sequence highlighted blue

**2) SpAAV-C intein (sequence with gRNA)**

TTGAGATCCTTTTTTTCTGCGCGTAATCTGCTGCTTGCAAACAAAAAAACCACCGCTA
CCAGCGGTGGTTTGTTTGCCGGATCAAGAGCTACCAACTCTTTTTCCGAAGGTAACTG
GCTTCAGCAGAGCGCAGATACCAAATACTGTTCTTCTAGTGTAGCCGTAGTTAGGCCA
CCACTTCAAGAACTCTGTAGCACCGCCTACATACCTCGCTCTGCTAATCCTGTTACCA
GTGGCTGCTGCCAGTGGCGATAAGTCGTGTCTTACCGGGTTGGACTCAAGACGATAG
TTACCGGATAAGGCGCAGCGGTCGGGCTGAACGGGGGGTTCGTGCACACAGCCCAG
CTTGGAGCGAACGACCTACACCGAACTGAGATACCTACAGCGTGAGCTATGAGAAAG
CGCCACGCTTCCCGAAGGGAGAAAGGCGGACAGGTATCCGGTAAGCGGCAGGGTCG
GAACAGGAGAGCGCACGAGGGAGCTTCCAGGGGGAAACGCCTGGTATCTTTATAGTC
CTGTCGGGTTTCGCCACCTCTGACTTGAGCGTCGATTTTTGTGATGCTCGTCAGGGGG
GCGGAGCCTATGGAAAAACGCCAGCAACGCGGCCTTTTTACGGTTCCTGGCCTTTTG
CTGGCCTTTTGCTCACATGTTCTTTCCTGCGTTATCCCCTGATTCTGTGGATAACCGTAT
TACCGCCTTTGAGTGAGCTGATACCGCTCGCCGCAGCCGAACGACCGAGCGCAGCGA
GTCAGTGAGCGAGGAAGCGGAAGAGCGCCCAATACGCAAACCGCCTCTCCCCGCGC
GTTGGCCGATTCATTAATGCAGCTGGCACGACAGGTTTCCCGACTGGAAAGCGGGCA
GTGAGCGCAACGCAATTAATGTGAGTTAGCTCACTCATTAGGCACCCCAGGCTTTACA
CTTTATGCTTCCGGCTCGTATGTTGTGTGGAATTGTGAGCGGATAACAATTTCACACAG
GAAACAGCTATGACCATGATTACGCCAGATTTAATTAAGGCTGCGCGCTCGCTCGCTC
ACTGAGGCCGCCCGGGCAAAGCCCGGGCGTCGGGCGACCTTTGGTCGCCCGGCCTC
AGTGAGCGAGCGAGCGCGCAGAGAGGGAGTGGCCAACTCCATCACTAGGGGTTCCT
GCGGCCTCTAGATCAGGGTACCCGTTACATAACTTACGGTAAATGGCCCGCCTGGCTG
ACCGCCCAACGACCCCCGCCCATTGACGTCAATAGTAACGCCAATAGGGACTTTCCAT
TGACGTCAATGGGTGGAGTATTTACGGTAAACTGCCCACTTGGCAGTACATCAAGTGT
ATCATATGCCAAGTACGCCCCCTATTGACGTCAATGACGGTAAATGGCCCGCCTGGCA
TTGTGCCCAGTACATGACCTTATGGGACTTTCCTACTTGGCAGTACATCTACGTATTAG
TCATCGCTATTACCATGGTCGAGGTGAGCCCCACGTTCTGCTTCACTCTCCCCATCTCC
CCCCCCTCCCCACCCCCAATTTTGTATTTATTTATTTTTTAATTATTTTGTGCAGCGATG
GGGGCGGGGGGGGGGGGGGCGCGCGCCAGGCGGGGCGGGGCGGGGCGAGGGG
CGGGGCGGGGCGAGGCGGAGAGGTGCGGCGGCAGCCAATCAGAGCGGCGCGCTCC

GAAAGTTTCCTTTTATGGCGAGGCGGCGGCGGCGGCCCTATAAAAAGCGAAGCG
CGCGGCGGGCGGGAGTCGCTGCGCGCTGCCTTCGCCCCGTGCCCCGCTCCGCCGCCG
CCTCGCGCCGCCCGCCCCGGCTCTGACTGACCGCGTTACTCCCACAGGTGAGCGGGC
GGGACGGCCCTTCTCCTCCGGGCTGTAATTAGCTGAGCAAGAGGTAAGGGTTTAAGG
GATGGTTGGTTGGTGGGGTATTAATGTTTAATTACCTGGAGCACCTGCCTGAAATCAC
TTTTTTTCAGGTTGGACCGGTGCCACCATGAAACGGACAGCCGACGGAAGCGAGTTC
GAGTCACCAAAGAAGAAGCGGAAAGTCATCAAGATTGCTACACGGAAATACCTGGG
AAAGCAGAACGTGTACGACATCGGCGTGGAGCGGGATCACAACTTCGCCCTGAAGA
ATGGCTTTATCGCCAGCAATTGCTTCGACTCCGTGGAAATCTCCGGCGTGGAAGATCG
GTTCAACGCCTCCCTGGGCACATACCACGATCTGCTGAAAATTATCAAGGACAAGGA
CTTCCTGGACAATGAGGAAAACGAGGACATTCTGGAAGATATCGTGCTGACCCTGAC
ACTGTTTGAGGACAGAGAGATGATCGAGGAACGGCTGAAAACCTATGCCCACCTGTT
CGACGACAAAGTGATGAAGCAGCTGAAGCGGCGGAGATACACCGGCTGGGGCAGGC
TGAGCCGGAAGCTGATCAACGGCATCCGGGACAAGCAGTCCGGCAAGACAATCCTG
GATTTCCTGAAGTCCGACGGCTTCGCCAACAGAAACTTCATGCAGCTGATCCACGAC
GACAGCCTGACCTTTAAAGAGGACATCCAGAAAGCCCAGGTGTCCGGCCAGGGCGA
TAGCCTGCACGAGCACATTGCCAATCTGGCCGGCAGCCCCGCCATTAAGAAGGGCAT
CCTGCAGACAGTGAAGGTGGTGGACGAGCTCGTGAAAGTGATGGGCCGGCACAAGC
CCGAGAACATCGTGATCGAAATGGCCAGAGAGAACCAGACCACCCAGAAGGGACAG
AAGAACAGCCGCGAGAGAATGAAGCGGATCGAAGAGGGCATCAAAGAGCTGGGCA
GCCAGATCCTGAAAGAACACCCCGTGGAAAACACCCAGCTGCAGAACGAGAAGCTG
TACCTGTACTACCTGCAGAATGGGCGGGATATGTACGTGGACCAGGAACTGGACATCA
ACCGGCTGTCCGACTACGATGTGGACCATATCGTGCCTCAGAGCTTTCTGAAGGACG
ACTCCATCGACAACAAGGTGCTGACCAGAAGCGACAAGAACCGGGGCAAGAGCGA
CAACGTGCCCTCCGAAGAGGTCGTGAAGAAGATGAAGAACTACTGGCGGCAGCTGC
TGAACGCCAAGCTGATTACCCAGAGAAAGTTCGACAATCTGACCAAGGCCGAGAGA
GGCGGCCTGAGCGAACTGGATAAGGCCGGCTTCATCAAGAGACAGCTGGTGGAAAC
CCGGCAGATCACAAAGCACGTGGCACAGATCCTGGACTCCCGGATGAACACTAAGTA
CGACGAGAATGACAAGCTGATCCGGGAAGTGAAAGTGATCACCCTGAAGTCCAAGC
TGGTGTCCGATTTCCGGAAGGATTTCCAGTTTTACAAAGTGCGCGAGATCAACAACTA
CCACCACGCCCACGACGCCTACCTGAACGCCGTCGTGGGAACCGCCCTGATCAAAAA

GTACCCTAAGCTGGAAAGCGAGTTCGTGTACGGCGACTACAAGGTGTACGACGTGCG
GAAGATGATCGCCAAGAGCGAGCAGGAAATCGGCAAGGCTACCGCCAAGTACTTCTT
CTACAGCAACATCATGAACTTTTTCAAGACCGAGATTACCCTGGCCAACGGCGAGAT
CCGGAAGCGGCCTCTGATCGAGACAAACGGCGAAACCGGGGAGATCGTGTGGGATA
AGGGCCGGGATTTTGCCACCGTGCGGAAAGTGCTGAGCATGCCCCAAGTGAATATCG
TGAAAAAGACCGAGGTGCAGACAGGCGGCTTCAGCAAAGAGTCTATCCTGCCCAAG
AGGAACAGCGATAAGCTGATCGCCAGAAAGAAGGACTGGGACCCTAAGAAGTACGG
CGGCTTCGACAGCCCCACCGTGGCCTATTCTGTGCTGGTGGTGGCCAAAGTGGAAAA
GGGCAAGTCCAAGAAACTGAAGAGTGTGAAAGAGCTGCTGGGGATCACCATCATGG
AAAGAAGCAGCTTCGAGAAGAATCCCATCGACTTTCTGGAAGCCAAGGGCTACAAA
GAAGTGAAAAAGGACCTGATCATCAAGCTGCCTAAGTACTCCCTGTTCGAGCTGGAA
AACGGCCGGAAGAGAATGCTGGCCTCTGCCGGCGAACTGCAGAAGGGAAACGAACT
GGCCCTGCCCTCCAAATATGTGAACTTCCTGTACCTGGCCAGCCACTATGAGAAGCTG
AAGGGCTCCCCCGAGGATAATGAGCAGAAACAGCTGTTTGTGGAACAGCACAAGCA
CTACCTGGACGAGATCATCGAGCAGATCAGCGAGTTCTCCAAGAGAGTGATCCTGGC
CGACGCTAATCTGGACAAAGTGCTGTCCGCCTACAACAAGCACCGGGATAAGCCCAT
CAGAGAGCAGGCCGAGAATATCATCCACCTGTTTACCCTGACCAATCTGGGAGCCCC
TGCCGCCTTCAAGTACTTTGACACCACCATCGACCGGAAGAGGTACACCAGCACCAA
AGAGGTGCTGGACGCCACCCTGATCCACCAGAGCATCACCGGCCTGTACGAGACACG
GATCGACCTGTCTCAGCTGGGAGGTGACAGCGGCGGGAGCGGCGGGAGCGGGGGG
AGCACTAATCTGAGCGACATCATTGAGAAGGAGACTGGGAAACAGCTGGTCATTCAG
GAGTCCATCCTGATGCTGCCTGAGGAGGTGGAGGAAGTGATCGGCAACAAGCCAGA
GTCTGACATCCTGGTGCACACCGCCTACGACGAGTCCACAGATGAGAATGTGATGCT
GCTGACCTCTGACGCCCCCGAGTATAAGCCTTGGGCCCTGGTCATCCAGGATTCTAAC
GGCGAGAATAAGATCAAGATGCTGAGCGGAGGATCCAAAAGAACCGCCGACGGCAG
CGAATTCGAGCCCAAGAAGAAGAGGAAAGTCTAAGATCTGATAATCAACCTCTGGAT
TACAAAATTTGTGAAAGATTGACTGGTATTCTTAACTATGTTGCTCCTTTTACGCTATG
TGGATACGCTGCTTTAATGCCTTTGTATCATGCTATTGCTTCCCGTATGGCTTTCATTTT
CTCCTCCTTGTATAAATCCTGGTTAGTTCTTGCCACGGCGGAACTCATCGCCGCCTGCC
TTGCCCGCTGCTGGACAGGGGCTCGGCTGTTGGGCACTGACAATTCCGTGGTGCGAC
TGTGCCTTCTAGTTGCCAGCCATCTGTTGTTTGCCCCTCCCCCGTGCCTTCCTTGACCC

TGGAAGGTGCCACTCCCACTGTCCTTTCCTAATAAAATGAGGAAATTGCATCGCATTG
TCTGAGTAGGTGTCATTCTATTCTGGGGGGTGGGGTGGGGCAGGACAGCAAGGGGGA
GGATTGGGAAGACAATAGCAGGCATGCTGGGGATGCGGTGGGCTCTATGGCAAAAAA
AGCACCGACTCGGTGCCACTTTTTCAAGTTGATAACGGACTAGCCTTATTTTAACTTG
CTATTTCTAGCTCTAAAACGCAGTTCTTCTTCTGTGTTGCGGTGTTTCGTCCTTTCCAC
AAGATATATAAAGCCAAGAAATCGAAATACTTTCAAGTTACGATAAGCATATGATAGTC
CATTTTAAAACATAATTTTAAAACTGCAAACTACCCAAGAAATTATTACTTTCTACGTC
ACGTATTTTGTACTAATATCTTTGTGTTTACAGTCAAATTAATTCCAATTATCTCTCTAA
CAGCCTTGTATCGTATATGCAAATATGAAGGAATCATGGGAAATAGGCCCTCGCAGGA
ACCCCTAGTGATGGAGTTGGCCACTCCCTCTCTGCGCGCTCGCTCGCTCACTGAGGCC
GGGCGACCAAAGGTCGCCCGACGCCCGGGCTTTGCCCGGGCGGCCTCAGTGAGCGA
GCGAGCGCGCAGCCTTAATTAACCTAATTCACTGGCCGTCGTTTTACAACGTCGTGAC
TGGGAAAACCCTGGCGTTACCCAACTTAATCGCCTTGCAGCACATCCCCCTTTCGCCA
GCTGGCGTAATAGCGAAGAGGCCCGCACCGATCGCCCTTCCCAACAGTTGCGCAGCC
TGAATGGCGAATGGGACGCGCCCTGTAGCGGCGCATTAAGCGCGGCGGGTGTGGTGG
TTACGCGCAGCGTGACCGCTACACTTGCCAGCGCCCTAGCGCCCGCTCCTTTCGCTTT
CTTCCCTTCCTTTCTCGCCACGTTCGCCGGCTTTCCCCGTCAAGCTCTAAATCGGGGG
CTCCCTTTAGGGTTCCGATTTAGTGCTTTACGGCACCTCGACCCCAAAAAACTTGATT
AGGGTGATGGTTCACGTAGTGGGCCATCGCCCTGATAGACGGTTTTTCGCCCTTTGAC
GTTGGAGTCCACGTTCTTTAATAGTGGACTCTTGTTCCAAACTGGAACAACACTCAAC
CCTATCTCGGTCTATTCTTTTGATTTATAAGGGATTTTGCCGATTTCGGCCTATTGGTTA
AAAAATGAGCTGATTTAACAAAAATTTAACGCGAATTTTAACAAAATATTAACGCTTA
CAATTTAGGTGGCACTTTTCGGGGAAATGTGCGCGGAACCCCTATTTGTTTATTTTTCT
AAATACATTCAAATATGTATCCGCTCATGAGACAATAACCCTGATAAATGCTTCAATAA
TATTGAAAAAGGAAGAGTATGAGTATTCAACATTTCCGTGTCGCCCTTATTCCCTTTTT
TGCGGCATTTTGCCTTCCTGTTTTTGCTCACCCAGAAACGCTGGTGAAAGTAAAAGAT
GCTGAAGATCAGTTGGGTGCACGAGTGGGTTACATCGAACTGGATCTCAACAGCGGT
AAGATCCTTGAGAGTTTTCGCCCCGAAGAACGTTTTCCAATGATGAGCACTTTTAAAG
TTCTGCTATGTGGCGCGGTATTATCCCGTATTGACGCCGGGCAAGAGCAACTCGGTCG
CCGCATACACTATTCTCAGAATGACTTGGTTGAGTACTCACCAGTCACAGAAAAGCAT
CTTACGGATGGCATGACAGTAAGAGAATTATGCAGTGCTGCCATAACCATGAGTGATA

ACACTGCGGCCAACTTACTTCTGACAACGATCGGAGGACCGAAGGAGCTAACCGCTT
TTTTGCACAACATGGGGGATCATGTAACTCGCCTTGATCGTTGGGAACCGGAGCTGA
ATGAAGCCATACCAAACGACGAGCGTGACACCACGATGCCTGTAGCAATGGCAACAA
CGTTGCGCAAACTATTAACTGGCGAACTACTTACTCTAGCTTCCCGGCAACAATTAAT
AGACTGGATGGAGGCGGATAAAGTTGCAGGACCACTTCTGCGCTCGGCCCTTCCGGC
TGGCTGGTTTATTGCTGATAAATCTGGAGCCGGTGAGCGTGGGTCTCGCGGTATCATT
GCAGCACTGGGGCCAGATGGTAAGCCCTCCCGTATCGTAGTTATCTACACGACGGGG
AGTCAGGCAACTATGGATGAACGAAATAGACAGATCGCTGAGATAGGTGCCTCACTG
ATTAAGCATTGGTAACTGTCAGACCAAGTTTACTCATATATACTTTAGATTGATTTAAA
ACTTCATTTTTAATTTAAAAGGATCTAGGTGAAGATCCTTTTTGATAATCTCATGACCA
AAATCCCTTAACGTGAGTTTTCGTTCCACTGAGCGTCAGACCCCGTAGAAAAGATCA
AAGGAT

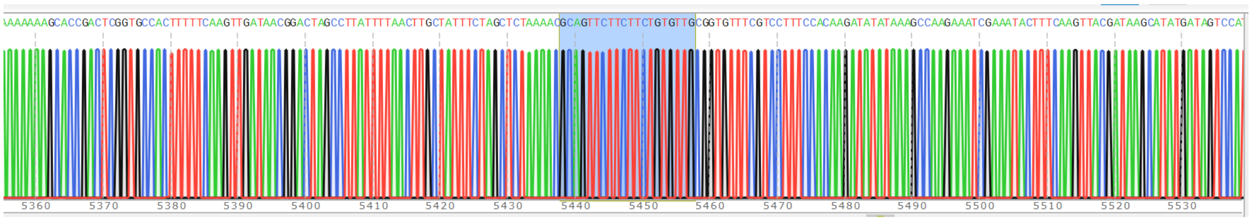


Sanger trace file – gRNA sequence highlighted blue

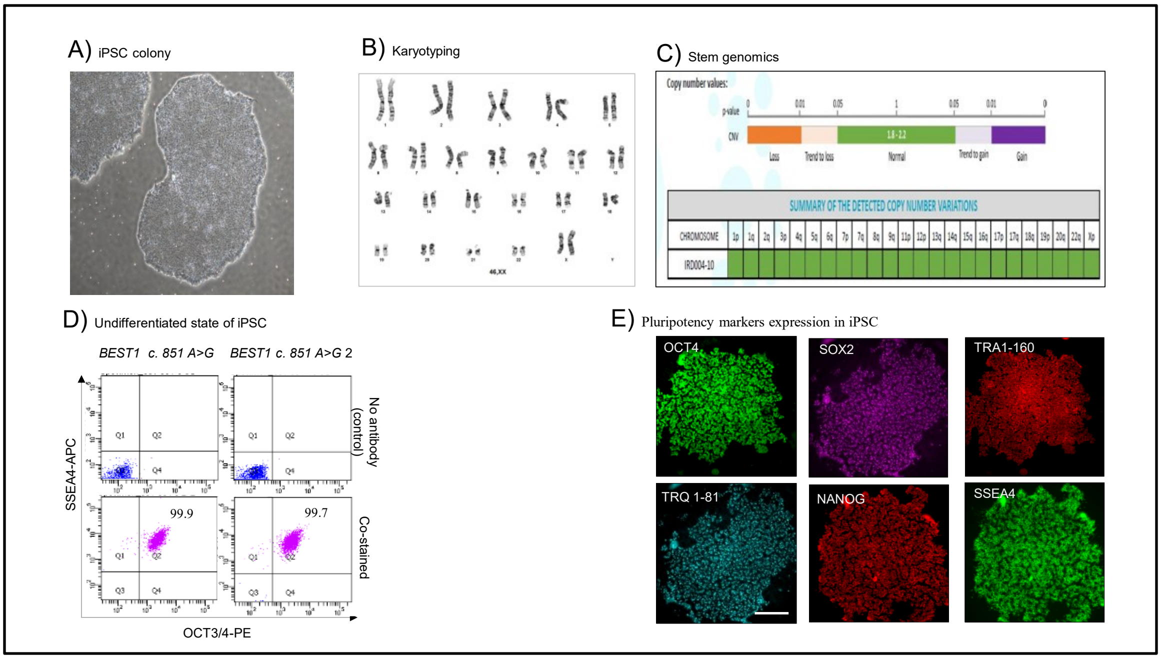


Figure S1: BEST1 variant iPSC characterization. (A) Morphological representation of a mature iPS colony using the Nikon TS100 Eclipse and 4x magnification (B)Characteristically normal karyotyping of an iPS cell line completed by the Cytogenetics Core at CHLA (C)Copy number variance performed by Stem Genomics (D) Flow cytometric analysis of undifferentiated state of iPSC markers OCT3/4 and SSEA4 showing >99% positive cell population, confirming robust pluripotency (E) Immunocytochemistry demonstrated the expression of pluripotency markers (SOX2,OCT4, SSEA4, NANOG,TRA1-160 and TRA-1-81) in iPSC lines of BEST1 c.851 A>G subjects (scale bar = 50 µm).

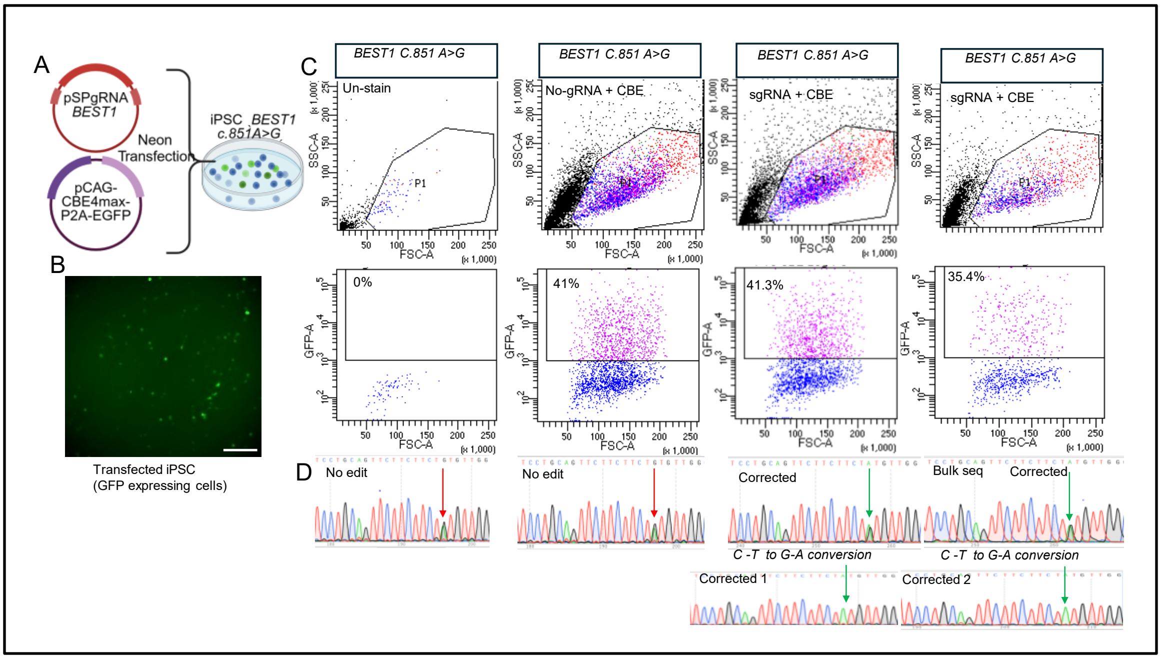


**Figure S2.** Generation of an isogenic control for iPSC line derived from *BEST1 C.851 A>G* patient blood (A) iPSCs were transfected with a base editor with GFP reporter (B) Representative fluorescent microscopy image showing GFP-positive iPSCs following transfection (scale bar = 50 µm) (C)Transfection efficiency was quantified by flow cytometry (FACS), with the dot plot indicating the percentage of GFP-positive cells (D) Bulk sequencing of edited cells confirmed successful genome editing, and both clonal populations and single-edited clones were subsequently isolated from the bulk-edited cells

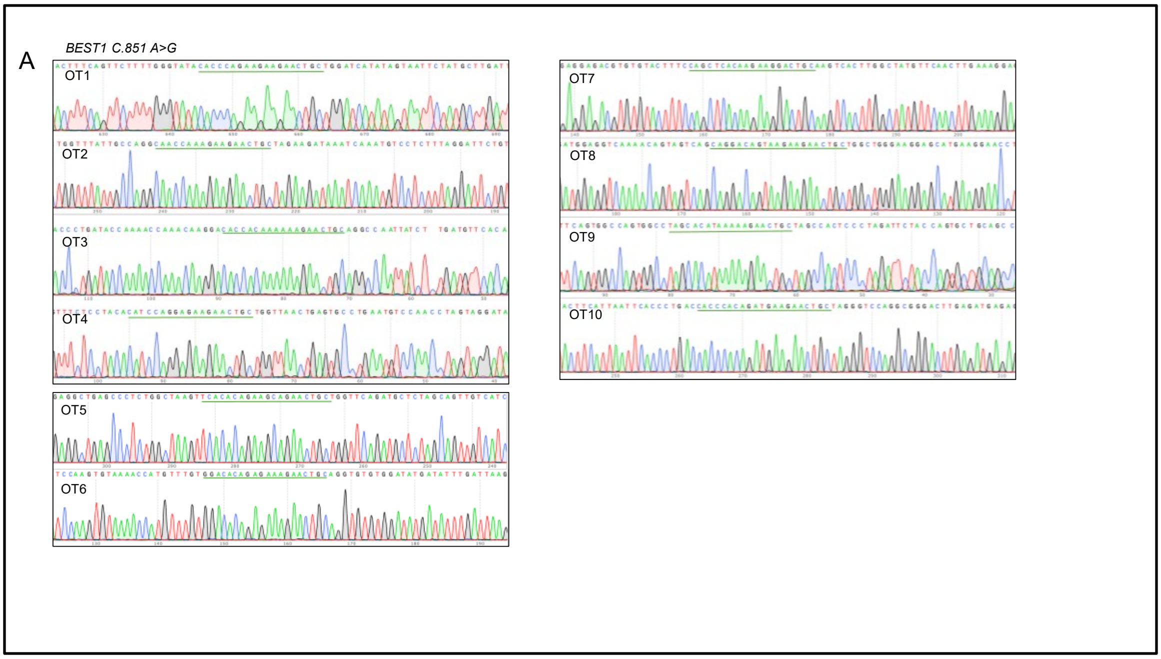


Figure S3- (A-C) Off-target validation in iPSCs with the BEST1 c.851A>G variant and two corrected clones derived from the parental line confirmed no off-target editing. (A) Off-target validation in iPSCs with the BEST1 c.851A>G variant , indicates no off target in notices in top ten predicted sites. OT- Off target. OT sequence marked by green line.

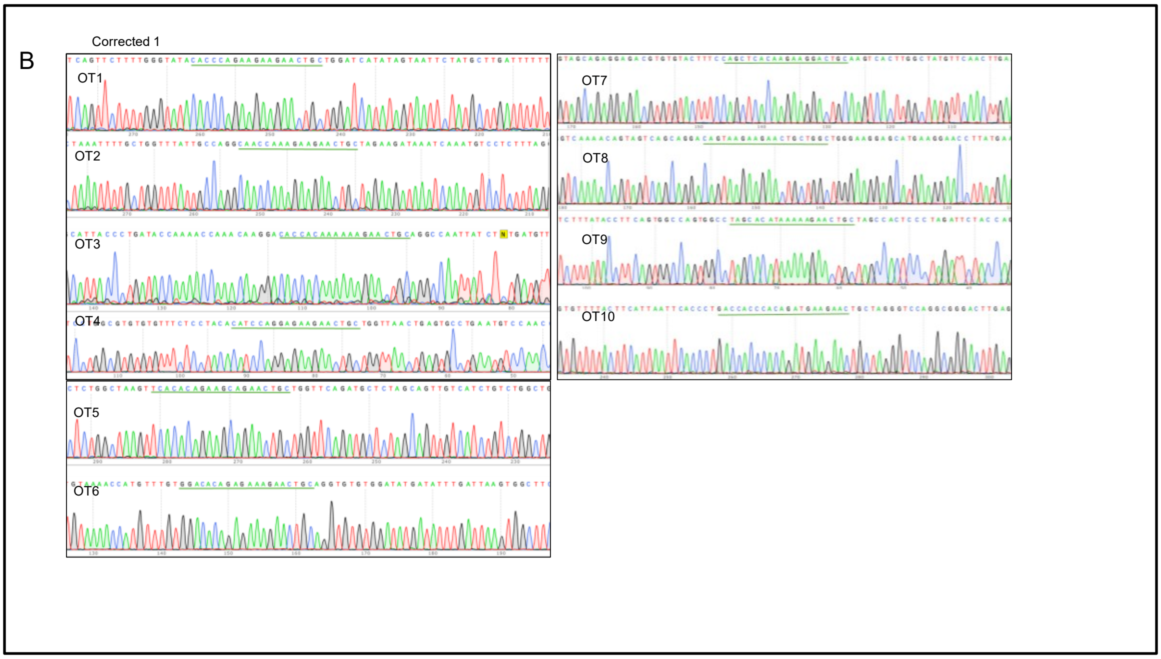


Figure S3 (B) Off-target validation in iPSCs with the corrected 1 indicates no off target in notices in top ten predicted sites

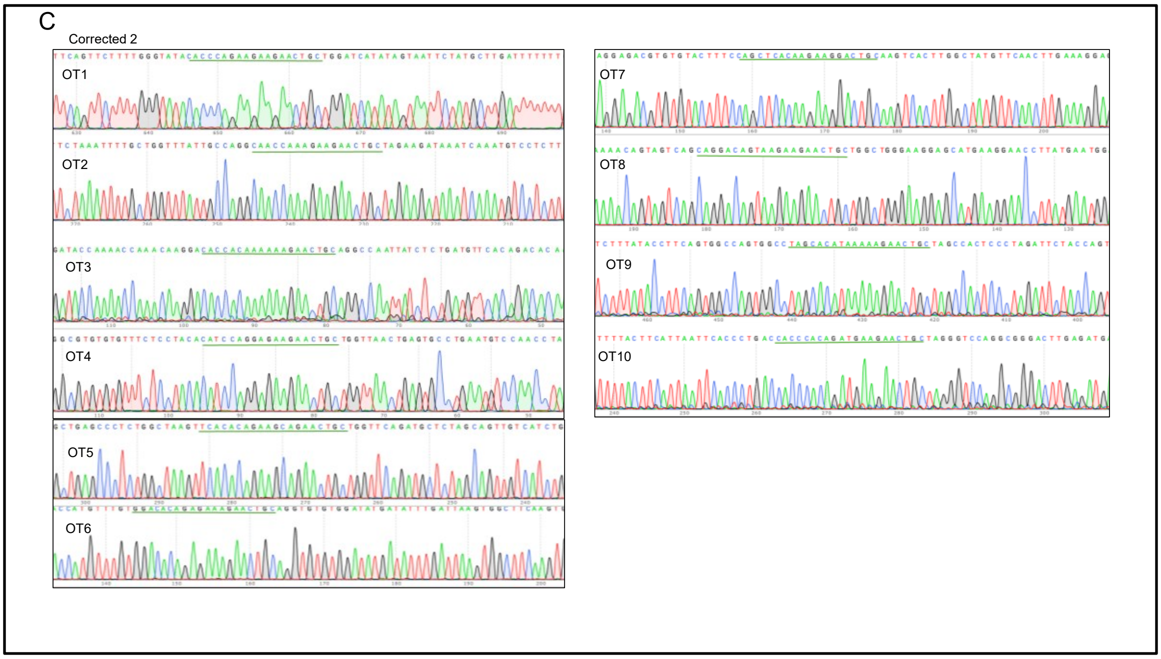


Figure S3 (C) Off-target validation in iPSCs with the corrected 2 indicates no off target in notices in top ten predicted sites

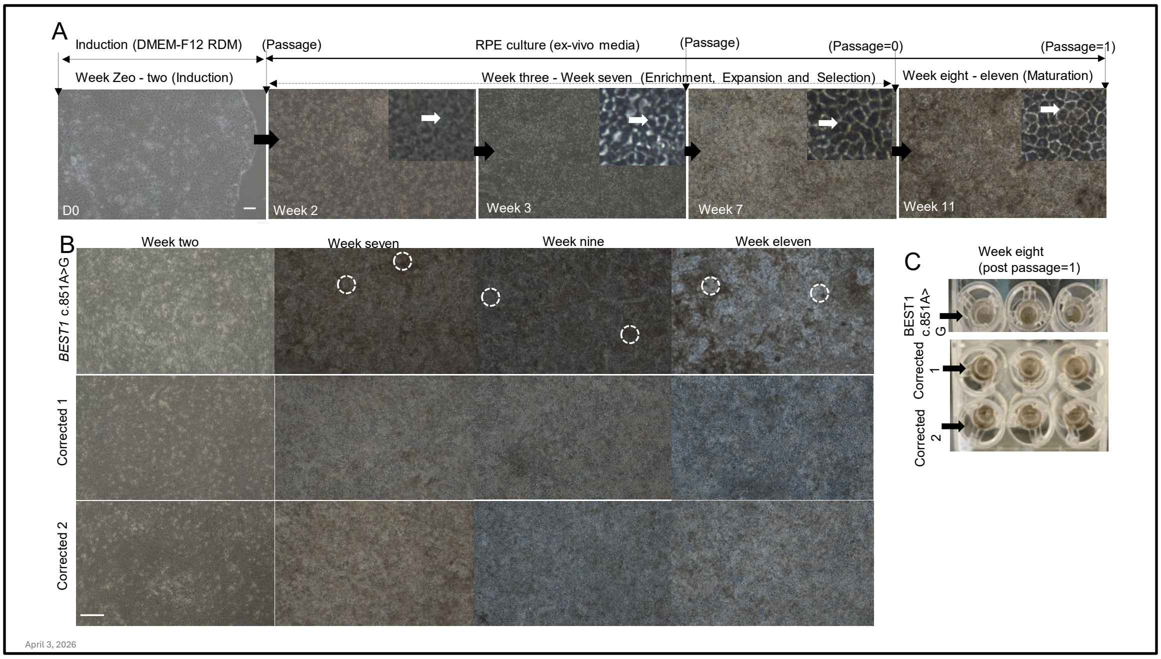


**Figure S4** RPE differentiation protocol and timeline. (A) RPE differentiation from iPSCs was carried out over 11 weeks. The protocol involved stepwise induction of retinal progenitors, followed by maturation into pigmented RPE cells. (B) Comparing *BEST1* mutant RPE and corrected RPE morphology. Mutant RPE shows focal loss (indicated by white dotted circle) at week 7-11 of culture, suggesting regions of necrotic cells. (C) RPE were cultured on transwell inserts, and images show pigmented RPE at week eight of differentiation (passage 1). Scale bar = 100µm . White Circle= Cell loss

**Induction phase media (Week 0-Week 2)**
**Day 0 and 1:** NIC, DKK, Noggin, IGF: anterior neuroectodermal/eye field induction
**Day 2**:NIC, DKK, Noggin, IGF, bFGF: neuroectodermal/eye field induction
**Day 4**: DKK, IGF, Activin A: anterior neuroectodermal/eye field and RPE induction
**Day 6**: Activin A, SU5402: RPE induction and retinal progenitor inhibition
**Day 8-14**: Activin A, SU5402, CHIR99021: RPE induction, retinal progenitor inhibition
**Enrich, Expansion, selection and maturation phase media ( Week 3-Week 11)**
**Day 14 –Day16** : *Ex-vivo* media, Penicillin and streptomycin, NIC, Rock inhibitor
**Day 19- beyond:** *Ex-vivo* media, Penicillin and streptomycin

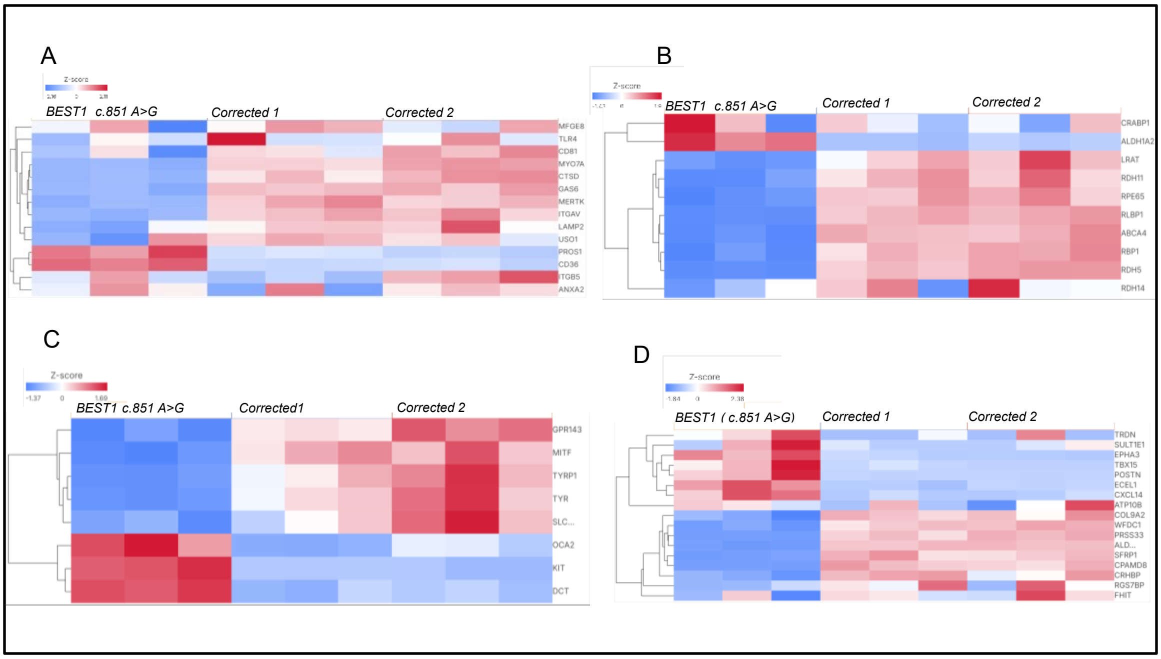


Figure S5-Transcriptomic profiling of BEST1 c.851 A>G variant and corrected clones reveals expression of RPE specific markers

Heatmaps showing transcriptional changes of RPE specific genes between BEST1 mutant and corrected clones. Heatmaps show transcriptional changes of RPE-specific genes from normalized expression data, categorized into (A) phagocytosis, (B) visual cycle, (C) pigmentation, and (D) macular RPE–associated genes.

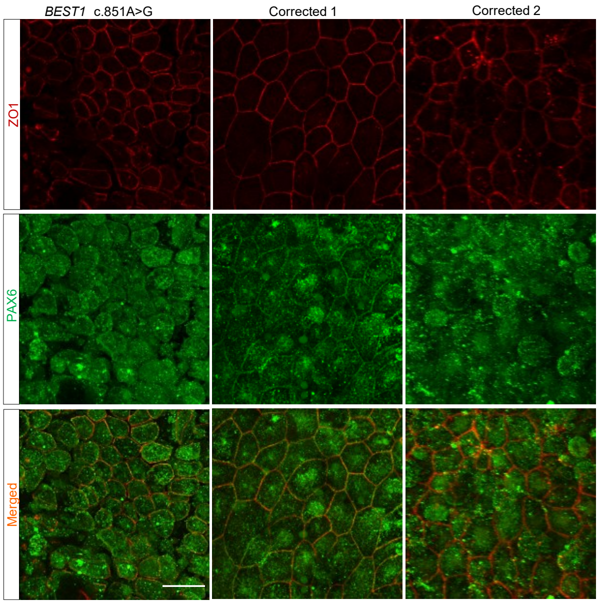


Figure S5. PAX6 expression in cultured RPE cells at week 4. Immunostaining shows nuclear PAX6 expression in RPE cells. ZO1 staining was used to select morphologically comparable RPE cells for analysis in BEST1 c.851A>G mutant and corrected lines 1 and 2. Scale bar= 20 µm

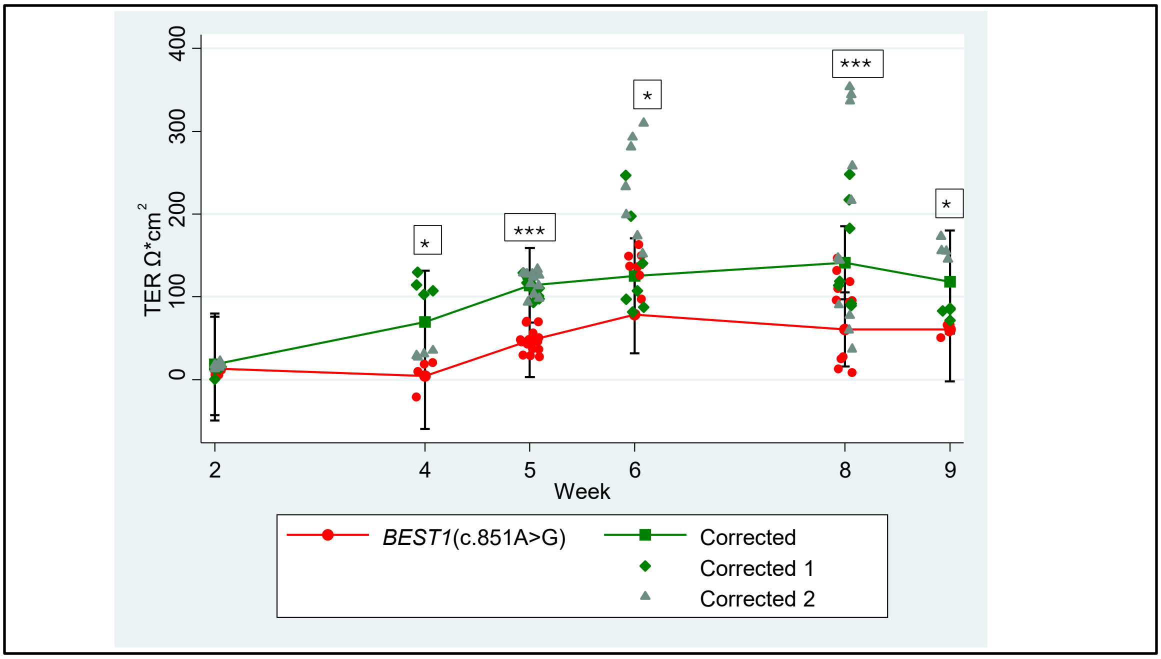


Figure S6- Transepithelial electrical resistance (TER) analysis over time. Individual data points represent replicates from weeks two-nine, showing differences in TER between BEST1 mutant and corrected RPE. Data are represented as mean ± SEM. Statistically significant differences in TER were observed between BEST1 c.851A>G mutant and corrected at all time points except week two (*= statistically significant).

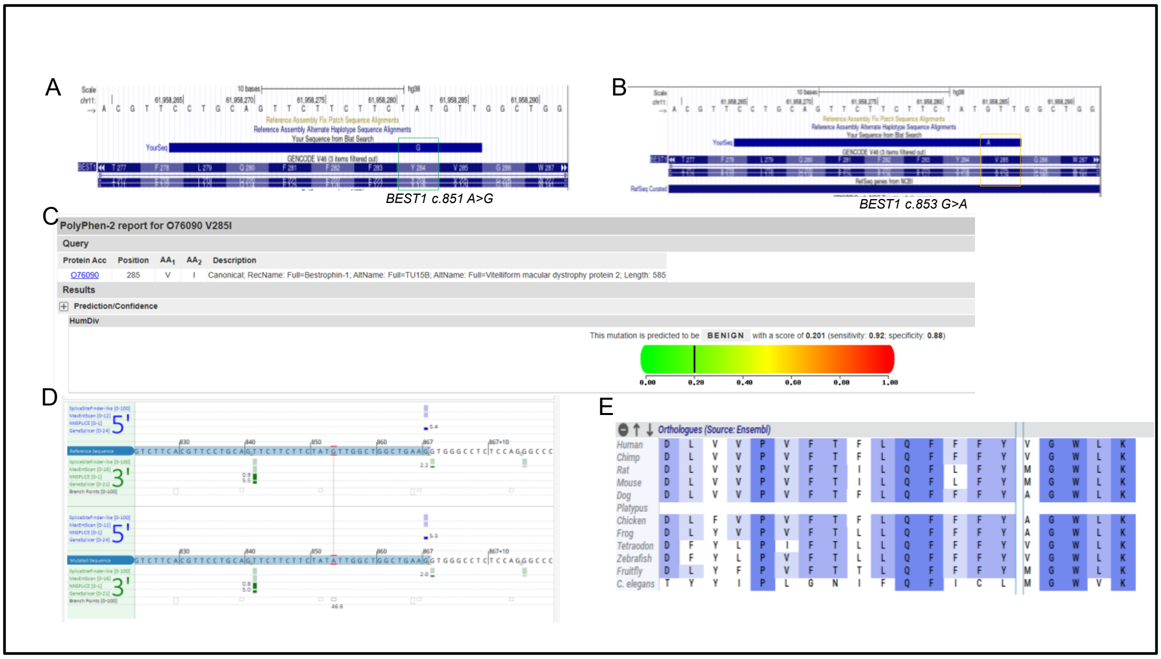


**Figure S7**: Integrative Genomics Viewer (IGV), showing a sequence alignment around the *BEST1* gene. *BEST1*(c.851A>G) bystander (c.853 G>A). Computation prediction. GTT (285)- Valine ATT (285)- Isoleucine ( bystander) as shown in (A) and (B). Valine is a hydrophobic amino acid which is commonly found in the interior of the proteins due to the presence of hydrophobic side chain. Valine has the second highest hydropathy index (4.2) of the polar amino acids, and its side chain is nonreactive and rarely participates in the protein function. (C-E) Computational prediction of the bystander edit indicates uncertain significance and low evolutionary conservation (4/12 compared species).
